# Phenome-scale causal network discovery with bidirectional mediated Mendelian randomization

**DOI:** 10.1101/2020.06.18.160176

**Authors:** Brielin C. Brown, David A. Knowles

## Abstract

Inference of directed biological networks from observational genomics datasets is a crucial but notoriously difficult challenge. Modern population-scale biobanks, containing simultaneous measurements of traits, biomarkers, and genetic variation, offer an unprecedented opportunity to study biological networks. Mendelian randomization (MR) has received attention as a class of methods for inferring causal effects in observational data that uses genetic variants as instrumental variables, but MR methods rely on assumptions that limit their application to complex traits at the biobank-scale. Moreover, MR estimates the total effect of one trait on another, which may be mediated by other factors. Biobanks include measurements of many potential mediators, in principle enabling the conversion of MR estimates into direct effects representing a causal network. Here, we show that this can be accomplished by a flexible two stage procedure we call *bidirectional mediated Mendelian randomization* (bimmer). First, bimmer estimates the effect of every trait on every other. Next, bimmer finds a parsimonious network that explains these effects using direct and mediated causal paths. We introduce novel methods for both steps and show via extensive simulations that bimmer is able to learn causal network structures even in the presence of non-causal genetic correlation. We apply bimmer to 405 phenotypes from the UK biobank and demonstrate that learning the network structure is invaluable for interpreting the results of phenome-wide MR, while lending causal support to several recent observational studies.

## 1 Introduction

Recent developments in the understanding of complex-trait genetics have lead to a call for increased study of directed biological networks because they are crucial for detecting core genes in the omnigenic model of complex traits, understanding risk factors for disease, and finding pathways that can be targeted for treatment [1, 2, 3]. However, interrogating the causal structure of networks is notoriously difficult owing to factors such as unmeasured confounding and reverse causation [4]. In spite of these challenges, modern population-scale biobanks offer an unprecedented opportunity to study biological networks because they contain measurements of traits, biomarkers, and genetic variation in the same individuals [5, 6]. Current methods for estimating biological networks from observational data with confounding are severely limited [7, 8, 9, 10] and have no strategy for integrating genetic data.

Mendelian randomization (MR) has recently received increased attention as a class of methods that can mitigate issues in causal inference by using genetic variants (SNPs) from genome-wide association studies (GWAS) as instrumental variables to determine the effect of an exposure (A) on an outcome (B). To estimate causal effects, MR methods must make strong assumptions that limit their ability to be applied at the biobank-scale. Perhaps the most controversial assumption is that the SNP only effects B through A (*i.e.* there is no horizontal pleiotropy). Recent methods such as Egger regression and the mode-based-estimator are able to relax this assumption, instead assuming there is no correlated pleiotropy or modal pleiotopy, respectively [11, 12]. Another approach, the latent causal variable (LCV) model, is able to detect causality under arbitrarily-structured pleiotropy [13]. However, the quantity that LCV calculates is not interpretable as the causal effect size of A on B. Most MR studies also presuppose the direction of effect, specifying one phenotype as the outcome and the other as the exposure. Pre-specifying the effect direction is sound when the outcome is clearly biologically downstream of the exposure, but in some cases it is better to learn the direction of the effect from the data. Some researchers have instead used bidirectional MR [14, 15], which tests for an effect in each direction, or gwas-pw [16], which infers the effect direction from the data. However, the utility of these approaches for complex traits, which might contain non-causal genetic correlation, is questionable [13].

In mimicking a randomized controlled trial, MR estimates the total causal effect (TCE) of A on B [17]. This effect may be mediated by any number of factors. The proliferation of phenome-scale datasets allows researchers to measure the effects of many possible mediators. In principle, this enables the conversion of TCE estimates into direct causal effect (DCE) estimates, which are not mediated by any other measured factor. However, methods to enact this conversion are limited, either because they require complex processing pipelines that limit their scope [18] or because they are computationally intractable for graphs with more than a few nodes [19]. This raises another disadvantage of approaches such as LCV and gwas-pw. Assuming that either A causes B or B causes A, but not both, is equivalent to assuming that the underlying causal network lacks cycles, which are thought to be an important part of real biological networks [20].

Here, we show that directed causal graphs can be estimated from phenome-scale GWAS summary statistics using a simple two-stage framework: first calculate the TCE of every phenotype on every other, then approximately invert that matrix to produce a causal graph. We call this framework *bi-directional mediated Mendelian randomization* (bimmer), and introduce novel methods for both components. To calculate the TCE matrix, we use a new weighting scheme for Egger regression that reduces the influence of pleiotropic SNPs (Figure 1a-b). Then, we convert the TCE estimates into a DCE graph via a novel algorithm for finding a sparse inverse to a partially-observed matrix that we call *inverse sparse regression* (inspre, Figure 1b-c). In extensive simulations, we show that our approach is able to learn causal network structures even in the presence of non-causal genetic correlation. We apply our method to 405 phenotypes from the UK Biobank, finding thousands of direct causal effects, complex causal pathways, and densely-connected sub-networks with correlated downstream effects.

**Figure 1:**
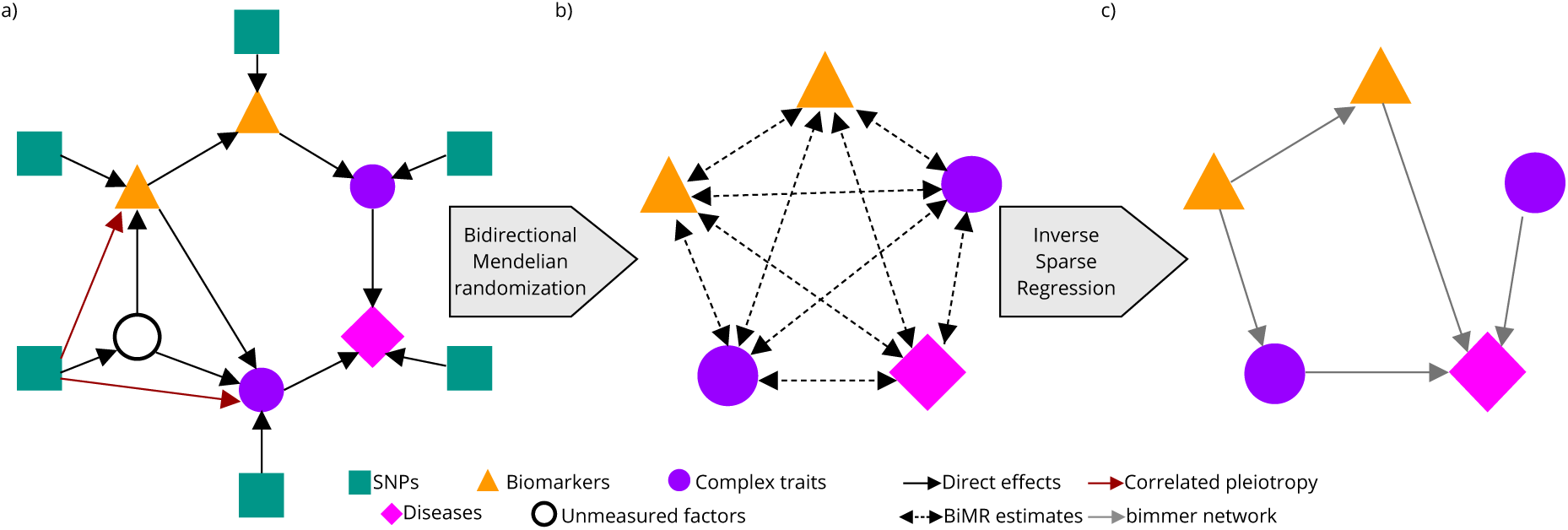
Overview of the bimmer model. a) Modern biobanks contains measurements of genetic variants (green squares), biomarkers (orange triangles), complex traits (purple circles) and diseases (pink diamonds). Genetic variants affect these phenotypes which in turn affect each other. The presence of unmeasured latent factors (white circle) can induce correlated pleiotropy (maroon arrows), but this effect can be reduced by down-weighting SNPs that seem to have an overly-strong effect on both phenotypes. b) Bi-directional Mendelian randomization estimates the total causal effect of the phenotypes on each other (dashed bi-directed arrows), which includes both direct and indirect effects. c) The direct effects can be found by estimating a sparse approximate inverse to the matrix of total effects, a process we call inverse sparse regression. This gives an estimate of the causal network (gray arrows), which can be imperfect. The modular nature of our approach allows us to use any method for bidirectional Mendelian randomization or approximate matrix inversion, providing flexibility and naturally accommodating future progress on both tasks.

## 2 Results

### Overview of model

We propose a simple regression model that nevertheless accommodates complex, bi-directional relationships. We assume each phenotype is a linear function of other phenotypes, genetic factors, and environmental factors. Assume we have *N* individuals, *D* phenotypes and *M* SNPs, with *Y* the matrix of phenotypes, *X* the genotype matrix, *β* the SNP effect matix and *γ* a matrix of unknown environmental effects. Let *G* be the *D* × *D* matrix of direct causal effects (the *causal graph*), with *G*_*i j*_ the DCE of phenotype *i* on phenotype *j*. We assume that phenotypes do not effect themselves (*G*_*i,i*_ = 0), and that the network is sparse (*G* has many entries that are 0). Our goal is to estimate *G* given summary statistics for the association of the genotypes *X* with the phenotypes *Y*. Our trait model is *Y* = *Y G* + *Xβ* + *γ*. Let *R* be the matrix of TCE estimates from MR, with *R*_*i,j*_ the total causal effect of phenotype *i* on phenotype *j* and *S*_*i,j*_ its standard error. We show in section 4 that under this model,

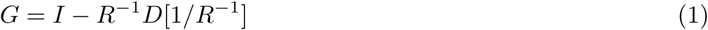

where *D* is an operator that sets all off-diagonal elements to 0, and */* represents element-wise division.

In practice, the matrix *R* need not be well-conditioned or even invertible, leading to challenges when calculating *G* via (1). Instead of calculating an exact or psuedo-inverse, we exploit the assumption that the underlying DCE matrix is sparse. Specifically, we seek matrices *U* and *V* such that *V U* = *I, U* ≈ *R* and *V* is sparse. We find them by solving the following constrained optimization problem,

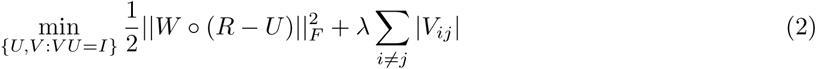

where 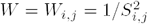 is a set of per-entry inverse variance weights, and *λ* is the *L*_1_ shrinkage parameter [21, 22]. We refer to *U* as bimmer shrunk estimates of the TCE, and use *V* ≈ *R*^−1^ to solve (1). Note that in this method, missing entries in *R* can be accommodated simply by setting their weights to 0. We use this property to our advantage in choosing the regularization parameter. Specifically, we use a novel adaptation of Stability Approach to Regularization Selection (StARS) [23] where we mask entries of the TCE in order to induce variance in the estimated graph during cross-validation. For complete details, see section 4.

Some intuition for (1) can be gained by considering the problem of estimating a matrix of partial correlations for a set of observed variables. Analogous to the DCE, the partial correlation measures the degree to which two variables are correlated while controlling for the effect of all other measured variables. Given a matrix of observed (standard) correlations, Σ, the matrix of partial correlations is *P* = −*D*[Σ^−1^]^−1*/*2^Σ^−1^*D*[Σ^−1^]^−1*/*2^. One of the most common approaches to obtaining a robust estimate of Σ^−1^, also called the precision matrix, is the graphical lasso (glasso) [21]. glasso assumes the data come from a multivariate normal distribution with a sparse precision matrix, and maximizes the likelihood with a *L*_1_ penalty on elements of Σ^−1^.

This leaves the problem of producing a reliable estimate for *R*, which can be particularly challenging when there is non-causal genetic correlation or differential power across phenotypes. Most MR studies use the set of genome-wide significant (GWS, *p*≤ 5 × 10^−8^) SNPs for a trait as instruments. Instead, we exploit the observation that in the absence of horizontal pleiotropy, if *A* causes *B* and a SNP effects *A* directly, the effect of the SNP on *B* can be no larger than the effect of the SNP on *A* times the effect of *A* on *B*. That is, the SNP must have its per-variance contribution to *B* reduced by the network. We use this intuition to construct a novel weighting scheme for Egger regression. First, we select a *p*-value threshold *p*_*t*_. For every phenotype *i*, we construct a set of marginally associated SNPs at threshold *p*_*t*_. Next, for every ordered pair of phenotypes *i, j*, we consider only SNPs that reach signficance level *p*_*t*_ in phenotype *i* but not *j*. For this set of SNPs, we calculate a weight based on the Welch test statistic for a two-sample difference in mean with unequal variances, and the standard inverse-variance weight. If 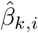 is our estimate of the effect of SNP *k* on phenotype *i* and *ŝ*_*k,i*_ its standard error, the Welch test statistic is [24]

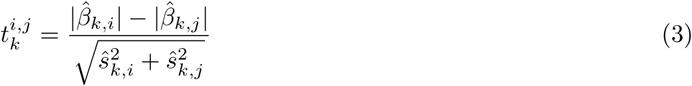

and our weight is 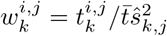. We use these SNP weights in the Egger regression of *j* on *i*. To avoid bias, we must use two sets of summary statistics. The first set is for SNP selection and weight construction, and the second set is for estimating *R*.

### Simulations

#### Weighted Egger regression improves calibration and power in Mendelian randomization

Our first goal was to assess whether our weighted Egger regression approach had a well-controlled type-I error rate (FPR) under the two-way null (no causal effect in either direction). To this end we simulated GWAS summary statistics for two phenotypes with *M* = 1, 000, 000 independent SNPs, 20% heritability and *N* = 100, 000 individuals in both the SNP discovery and effect estimation cohorts. In each simulation, there were 5, 000 causal SNPs per phenotype. In our first simulation, 1, 000 of these SNPs are pleiotropic, effecting both phenotypes, but with no correlation of their effects. In our second, these 1, 000 SNPs are again shared, but with equal effects on both phenotypes for a total genetic correlation of *ρ*_*g*_ = 0.2. In our final simulation under the null, we again have *ρ*_*g*_ = 0.2, except the phenotypes have very different sample sizes (*N*_1_ = 200, 000, *N*_2_ = 50, 000), and shared effects are twice as large on average for the phenotype with fewer samples. This makes shared SNPs much more likely to have low (significant) *p*-values in the second cohort. In each setting, we compared our approach against the standard approach of Egger regression using all SNPs reaching GWS for the exposure as instruments, as well as an oracle with access to the true effect sizes that uses only non-pleiotropic SNPs.

In the first setting, uncorrelated pleiotropy, all methods were able to effectively control the FPR at level *α* = 0.05 in both directions (Figure 2a, Table S1). In the second setting, correlated pleiotropy, standard Egger regression produced excess false-positives, but our weighting scheme is able to reduce the false positive rate substantially (Figure 2b, Table S1). In the most challenging setting, correlated pleiotropy with unequal power, standard Egger regression produces many excess false positives in both directions, but our weighting scheme again substantially reduces the error rate, from 0.284 to 0.087 in the *A*→ *B* direction and from 0.492 to 0.029 in the *B*→ *A* direction (Figure 2c, Table S1).

**Figure 2:**
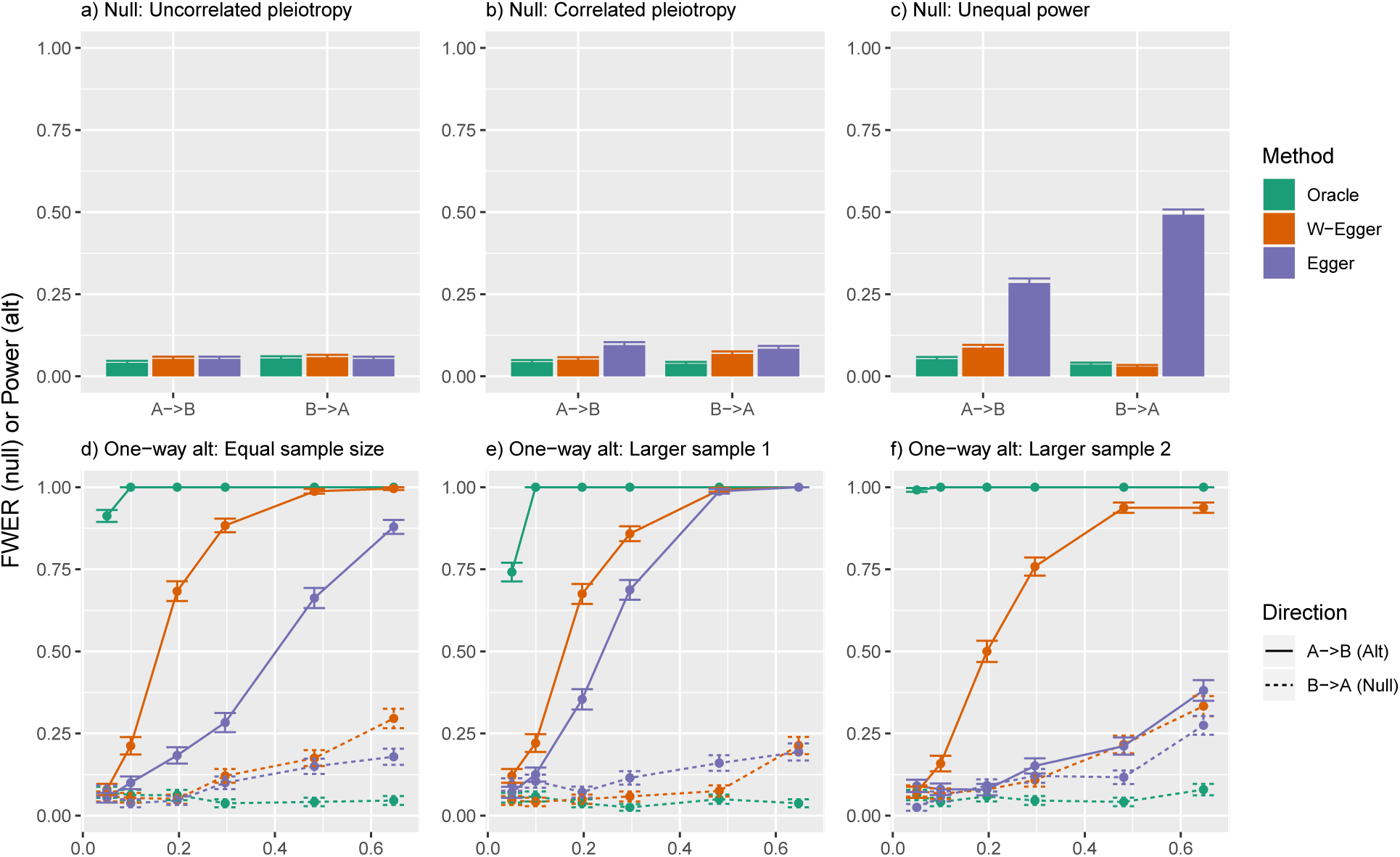
Weighted Egger regression reduces false positives and increases power in bi-directional MR. We simulated GWAS summary statistics for two phenotypes (*A, B*) with *M* = 1, 000, 000 independent SNPs, 20% heritability and *N* = 100, 000 individuals in both the SNP discovery and effect estimation cohorts. In each simulation, there were 5, 000 causal SNPs per phenotype. a) Both the effect of A on B and B on A are null, and 1000 of the SNPs have uncorrelated pleiotropic effects. All methods are well behaved. b) Both effects are again null, but the 1000 shared SNPs have equal effects on both phenotypes. Egger regression results in excess false positives which our weighting scheme reduces. c) Both effects are null and the shared SNPs have an equal effect on both phenotypes, but the shared SNPs have twice as large an effect on *B*, which also has a much smaller sample size. Egger regression results in numerous false positives, which our weighting scheme corrects. d) *A* has a variable effect on *B* and the studies have equal sample size. Our weighting scheme improves power over standard Egger regression. e) *A* effects *B*, which has a much lower sample size. Our weighting scheme improves power, but not as much as in (d). f) *A* effects *B*, but *A* has a much smaller sample size. Our weighting scheme substantially increases power. We conduced 1000 simulations for each null experiment (a-c) and 250 simulations per effect size for each alternative experiment (d-f).

Next, we wanted to asses the power of our approach under the one-way alternate hypothesis for various true effect sizes. We again conduct three simulations, calculating the power for effect sizes ranging from 0.05 to 0.7. In the first, the cohorts had equal sample sizes (*N* = 100, 000). In the second, the exposure cohort has larger sample size (*N*_1_ = 200, 000, *N*_2_ = 50, 000), and in the third the outcome cohort has a larger sample size (*N*_1_ = 50, 000, *N*_2_ = 200, 000). In all settings, our weighted Egger approach shows a substantial gain in power over standard Egger regression. This is especially notable for smaller effect sizes, and when the outcome GWAS is larger. In this latter setting, the power of standard Egger regression is only slightly higher than the FPR for the null hypothesis on the reverse direction, while our weighted Egger regression has very high power (Figure 2d-f, Table S2). However, both methods suffer from an increase in false positives in the reverse direction when the effect size in the forward direction is strong. For more on this phenomenon, see section 3.

Finally, we tested the power of our approach under the two-way alternate hypothesis. We tested pairs of effects ranging from −0.5 to 0.5 in both cohorts. Here we conduct two simulations: one with equal sample size of *N* = 100, 000, and one with unequal sample sizes *N*_1_ = 200, 000 and *N*_2_ = 50, 000. In all settings, our approach improves power substantially over standard Egger regression (Figure S1a-d). As with the one-way alternative, this is particularly apparent when the outcome has a larger sample size than the exposure (Figure S1d). We also observed that both methods had lower power when *R*_12_ ≈ −*R*_21_ and vice versa, especially when *R*_12_ has large absolute value. Indeed, as *R*_12_ →−*R*_21_→ 1, the model becomes unidentifiable. This setting is actually a violation of the *faithfulness* assumption commonly employed in causal inference [25].

In these simulations, we used a *p*-value threshold of 5× 10^−6^ for all weighted Egger regression analyses, but the conclusions held across a range from 5 ×10^−4^ to 5 ×10^−8^. We found that ×5 10^−6^ provided a reasonable balance between increased power under the alternative and control of type-I errors. However, lower cutoffs will provide better control of the type-I error rate in difficult situations at the expense of reduced power. Likewise, higher cut-offs yield higher power while reducing control of the type-I error rate (Table S3 and Table S4).

#### inspre is competitive with glasso while handling missing entries and directed graphs

As detailed above, both inspre and glasso can be viewed as methods for finding a sparse, approximate inverse to a noisily measured matrix. Therefore, we sought to compare these two methods with data simulated from an undirected graph with normally-distributed observations (the glasso model). We used the *huge* [26] package to generate data from a multivariate-normal distribution with a sparse precision matrix for various graph structures, sample sizes, and numbers of features. We considered three kinds of graph structures: 1) Erdös-Réyni (random) graphs, where each edge is included with probability *p*, 2) hub graphs, where nodes are partitioned into disjoint sets and every node in each set is connected to a central “hub” vertex, 3) scale-free graphs, where the vertex degree distribution follows a power law. Hub and scale-free networks are intended to mimic common biological networks [27]. We used the default edge weight in huge of ≈0.3. While it is important to evaluate both the presence of the edge and the accuracy of the inferred weight, the sparse nature of the problem renders traditional accuracy metrics such as the mean squared error uninformative. We follow standard convention [21, 23, 26] and focus on the precision, the number of true edges among all inferred edges, and recall, the proportion of true edges detected. We used these to calculate the *F*_1_ score, the harmonic mean of precision and recall, as a function of the stability of the inferred graph. For the graphical lasso, we used StARS to evaluate graph stability. For inspre, we used random masks in the weight matrix as detailed above.

First, we simulated data with 40 features and 800 samples. Our random graphs included each edge with probability *p* = 0.04, and our hub graphs had two hubs of 20 features each. In this setting inspre and glasso performed similarly for all graph types, with glasso performing slightly better on random graphs, inspre performing slightly better on hub graphs, and both methods having very similar performance for scale-free graphs (Figure S2a-c). Next, we simulated data with 100 features and 500 samples. Here our random graphs included each edge with probability *p* = 0.02 and our hub graphs had 5 hubs. In this setting, glasso outperformed inspre on random graphs and inspre outperformed glasso on hub graphs. Again both methods had similar performance on scale-free graphs, with a slight edge towards glasso (Figure S2d-f).

We hypothesized that if the entries in the correlation matrix had variable sample sizes, the ability of inspre to incorporate weights would improve performance relative to glasso. This represents a common real-world setting in which some features are measured on many samples, and some are measured on only a few. In each simulation, we first chose a maximum missingness threshold *m* uniformly between 50% and 99%. Then we simulated data with 100 features and 2000 samples. For each feature, we chose a number between 0 and *m* uniformly at random and set that proportion of the features samples as missing. We then calculated the sample correlation matrix using only samples where both features were measured per pair of features. In this setting, inspre was able to continue producing accurate results even when the maximum missingness was high. On the other hand, glasso was not able to produce results at all when there was high missingness. Instead, the glasso algorithm diverged and the program returned a matrix of NA values (Figure S3).

#### bimmer robustly recovers direct causal effect networks

Our final goal was to show that bi-directional Mendelian randomization could be combined with inspre to fit networks of simulated phenotypes from phenome-scale GWAS summary statistics. At the time of this writing we are not aware of any other methods for this specific problem. However, there are a few approaches to related problems that could be applied. Specifically, the DCEs between multiple exposures and a single outcome can be calculated from a multiple regression of SNP effects on the outcome against SNP effects on the exposures [28]. This approach can be used to find sparse effects by using a LASSO or elastic net regression (elnet-Egger). A more sophisticated approach, such as MR-Bayesian model averaging (MR-BMA), could also be applied [29].

First, we simulated summary statistics for 50 phenotypes with 1, 000 shared and 2, 000 private causal effect SNPs per pair of phenotypes: 125, 000 total SNPs. Each phenotype had 20% heritability. The causal network underlying the phenotypes came from an Erdös-Réyni random graph with randomly oriented edges. We found that MR-BMA performed comparably to elnet-Egger, but they both performed poorly compared to bimmer. Moreover, MR-BMA took about 20 times longer than bimmer to run with default parameter settings (Figure S4).

Next, we performed larger-scale simulations with 100 phenotypes and 250, 000 total SNPs. We again simulated data from Erdös-Réyni, hub, and scale-free networks. In this setting both the graph structure and the orientation of the graphs edges are important variables to consider. The edge orientation will not necessarily be random: for example, master regulators would have very high out-degree but low in-degree [23]. For all graph types, we tested three ways of orienting the edges in the graph: 1) randomly set the orientation of each edge (random), 2) preferentially orient edges towards high-degree nodes (towards), and 3) preferentially orient edges away from high-degree nodes (away). See Figure 3a-c for examples of different kinds of graphs with different edge orientations. We excluded MR-BMA from these simulations due to runtime concerns.

**Figure 3:**
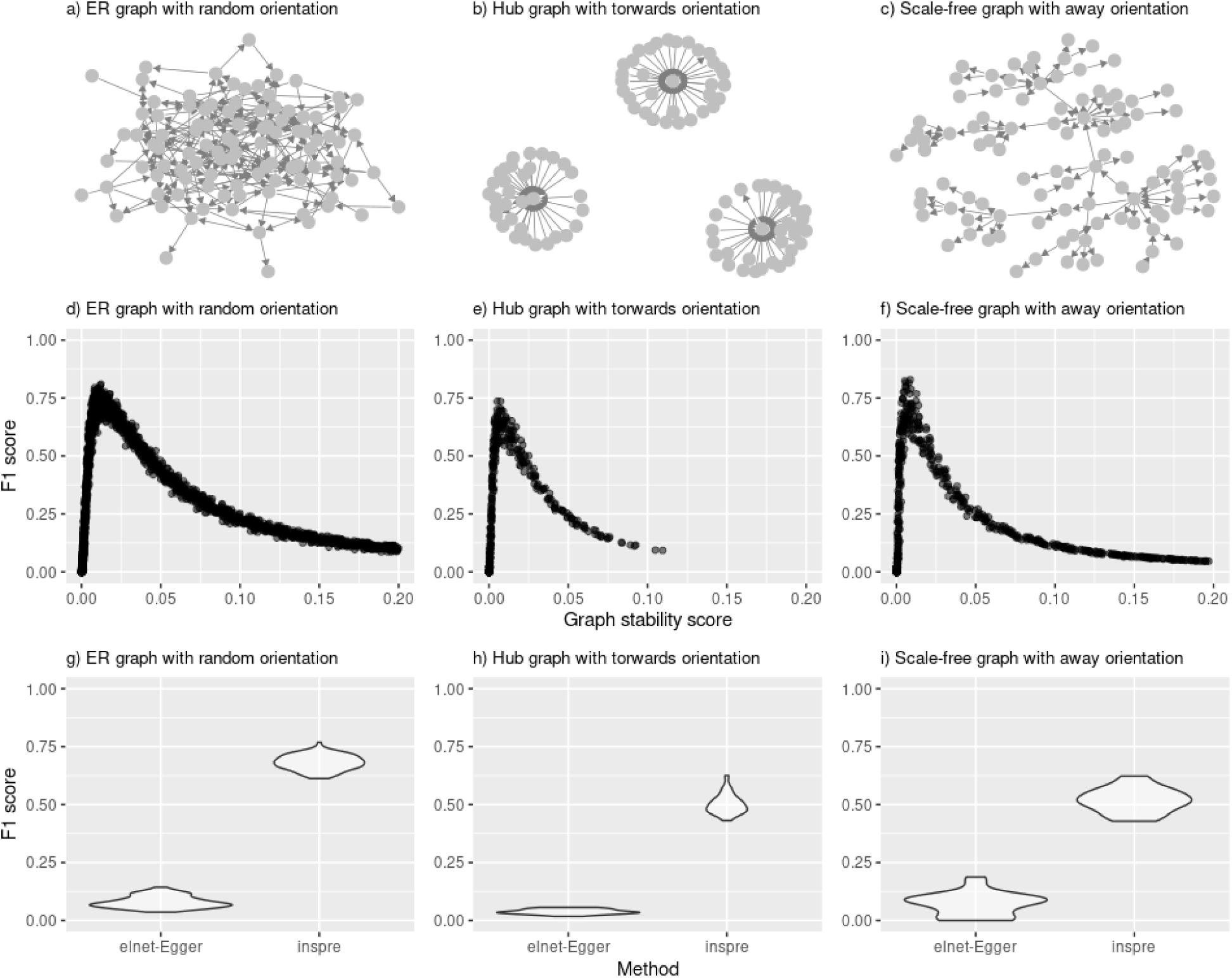
bimmer accurately infers the causal graph for many graph structures and node orientations. We simulated summary statistics for 100 phenotypes with 3000 causal effects each, 1000 of which were shared with uncorrelated effects per pair of phenotypes. We varied the structure and edge orientation of the causal graph underlying the phenotypes. a) An Erdos-Reyni random graph with randomly oriented edges. Each edge is included with probability *p* = 0.05 and then randomly assigned an orientation. b) A hub graph with edges preferentially oriented towards high degree nodes. The nodes are split into three sets and each node in each set is assigned to a central hub vertex. c) A scale-free graph with edges preferentially oriented away from high degree nodes. These graphs have node degrees that follow a power-law distribution. We show the *F*_1_-score of the method against the calculated variability score for d) Erdos-Reyni, e) hub and f) scale-free graphs. In all cases, we are able to produce accurate results when the variability score is between about 0.01 and 0.05. We also compared the performance against Egger regression with elastic-net shrinkage at a graph variability score of 0.025. elnet-Egger performs quite poorly compared to bimmer.

We found that bimmer was able to accurately re-construct all graph types and edge orientations considered, while elnet-Egger consistently had poor performance (Figure 3d-i, Figure S5). For Erdos-Reyni graphs, we found that edge orientation did not have an effect on the performance of bimmer. This is possibly because the node degree distribution does not have enough variance to have nodes that consistently pull edges towards or away from them in the latter scenarios. For scale-free and hub graphs, we found that bimmer performed better when high-degree nodes had edges oriented away from them (Figure 7). This is particularly interesting as it corresponds to the most likely real-world scenario [27]. Indeed, bimmer performed worst in the least realistic scenario: hub graphs with edges oriented towards the hubs (Figure 3e). However, even in this challenging setting bimmer is able to accurately infer the causal graph.

### Application to 405 traits from the UK Biobank

#### bimmer identifies thousands of direct causal effects in complex pathways

We obtained summary statistics for sex-split UK Biobank phenotypes from the Neale lab, who corrected for age, age^2^ and 20 principal components of the genotype matrix [30]. For ease of interpretation, we transformed all effect sizes to the per-variance scale. As previously suggested [31, 30], we used only phenotypes with *Z*-score above 4 and at least “medium” confidence. We removed one phenotype from every pair with genetic correlation above 0.9, leaving 423 phenotypes. We clumped the UKBB summary statistics to *p* = 5 × 10^−6^ with *r*^2^ < 0.05 and distance 500 kilobases using the UKBB European genotypes as a reference panel. We use male summary statistics for SNP selection and weight estimation, and female summary statistics for TCE estimation. Finally, we removed phenotypes where at least 50% of the standard errors of the TCE were above 0.5 as either an exposure or an outcome, resulting in 405 phenotypes (Table S5). 8, 268 (∼5%) of the 163, 620 pairs of traits considered had TCEs significant at FDR 5%.

We applied inspre (2) to the resulting TCE matrix (*R*) to infer the DCE network (*G*), using inverse-variance weights as previously described with one slight modification. To avoid having a very small number of pairs with very small SE dominate the loss, all entries in the TCE matrix with a standard error below 0.005 were given the same weight. We chose a target stability of 0.025 which gave reliable results across our various simulations (Figure 3, Figure S2, Figure S5). The resulting DCE graph had 7, 949 edges, which we pruned to 2, 826 edges by removing DCEs with absolute value less than 0.01.

We were curious to compare estimated genetic correlation, weighted Egger estimated TCE (*R*), bimmer’s shrunk TCE (*U*) and bimmer’s inferred DCE (*G*). First, we clustered phenotypes by genetic correlation to determine if the patterns observed are shared in the TCE estimates. While there are some similar patterns across the two matrices, the structure in the TCE estimates is not as well-defined (Figure S6a-b). Indeed, we find that while the TCE estimates and genetic correlation estimates are correlated, that correlation is fairly weak (*r* = 0.270 *±* 0.005). We actually find a slightly lower correlation between *U* and *R* (*r* = 0.238 *±* 0.005), but this is driven by TCE entries with high standard error that are consequently ignored by inspre’s weighting procedure. Restricting our analysis to TCE entries with an SE below 0.05, the correlation of the TCE with the genetic correlation is smaller than the correlation of the TCE with *U* (*r* = 0.666 *±* 0.005 vs *r* = 0.949 *±* 0.004, respectively). Moreover, entries in *U* tend to be close to 0 when the corresponding entry in the TCE has a large standard error (mean *|U|* = 0.0003 *±* 0.0003 for entries of the TCE with SE *>* 0.05). We conclude that bimmer produces a conservative estimate *U* ≈ *R* that accurately captures high confidence entries of *R* but performs aggressive shrinkage on edges with weak statistical support.

Most (303*/*405, ∼75%) phenotypes have out-degree 0 in our network, that is they have no downstream causal effects on the traits we consider. However, there is a path from every node with non-zero out-degree to every other node in the network. The majority of these connections are indirect and result in small effect sizes that do not reach statistical significance as TCEs; only 6, 007 of 39, 200 pairs of connected nodes have FDR-corrected TCE *p*-value below 0.05. Non-significant connections have an average absolute shrunk TCE of 0.0036 and a median DCE network path length of 3 nodes, while significant connections have an average shrunk TCE of 0.029 and median DCE network path length of 2 nodes (Figure 4a-b). Despite this, the majority (3, 831 connections, 63.7%) of significant TCEs result from indirect paths in the DCE network. The effect explained by the shortest path between two nodes is often only a small fraction of the total effect; there are typically multiple paths between two nodes that contribute to the TCE (Figure 4c-d). This finding is especially pronounced when looking at all connections, where the median percentage of TCE explained by shortest path is 0.35% (Figure 4c). For FDR 5%-significant connections, the median percentage of TCE explained by shortest path is 0.52% (Figure 4d). Interestingly, the percentage of the TCE explained by shortest path is sometimes greater than 1, i.e. the effect of the shortest path is greater than the total effect.

**Figure 4:**
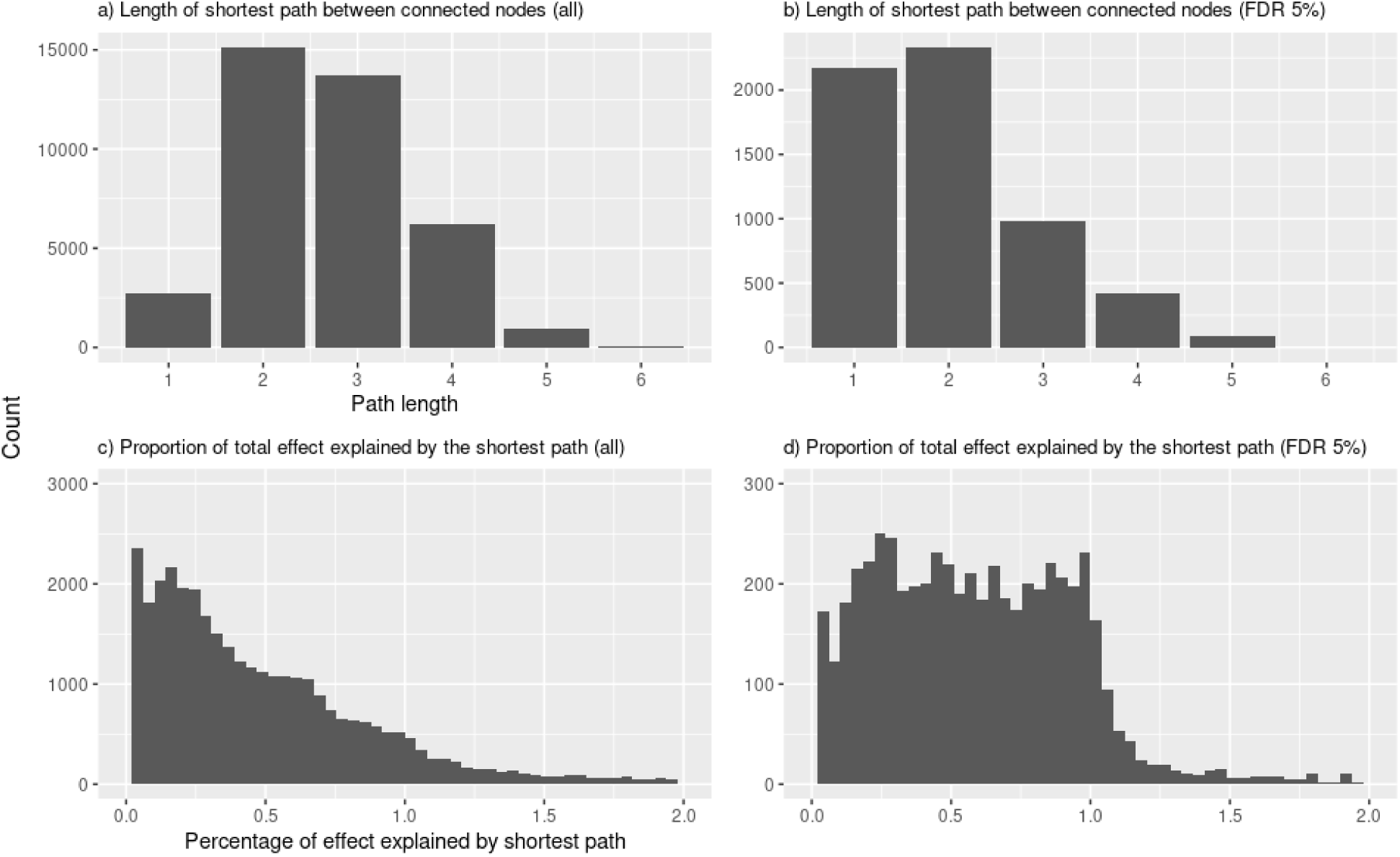
Applied to 405 phenotypes from the UK Biobank, bimmer identifies indirect effects and paths explaining a small proportion of the total effect. The distribution of path lengths between connected nodes for (a) all connected nodes and (b) connected nodes with an FDR 5% significant TCE. Our analysis shows that there are many long paths between nodes that result in very small effect sizes; closer connections are much more likely to reach significance after correction for multiple testing. Moreover the shortest path often explains only a fraction of the total effect for both (c) all connected nodes and (d) connected nodes with an FDR 5% significant TCE, indicating that there are often numerous ways of getting from one node to another that add together to form the TCE. We also observe that the shortest path sometimes explains more than the total effect, indicating that other paths between the nodes act to cancel out the effects of the direct path.

This occurs when other causal paths act to cancel out the effect of the shortest path and reduce the total effect.

Access to the DCE network, rather than just the TCEs, greatly aids data analysis and interpretation. For example, bimmer is able to impute the existence of some edges that have low statistical support as TCEs, but are required in order to explain other highly significant effects. Of the 2,826 edges in our network, 571 correspond to FDR-corrected TCE *p*-values above 0.05. A striking example of this is an effect of “time spent watching television” (TSWT) on body mass index (BMI). This has no initial statistical support (*R* = 0.008 *±* 0.1, *p* = 0.94) but bimmer infers a strong effect (*G* = 0.110) in order to explain strong effects of TSWT on numerous downstream phenotypes such as “usual walking pace” (*p* < 2 × 10^−5^, *U* =− 0.022), “wheezing in the chest” (*p* < 4 × 10^−5^, *U* = −0.018) and “father’s age at death” (*p* < 5 × 10^−4^, *U* = −0.010). This gives evidence of a direct causal effect of a sedentary lifestyle on higher BMI, contradicting an earlier study using bi-directional Mendelian randomization that found a causal effect of higher BMI on less exercise [15]. We also observed a strong TCE of “age first had sexual intercourse” (AFSI) on a number of surprising outcomes including knee pain (*U* = −0.064, *p* < 3 × 10^−7^), wheezing in the chest (*U* = −0.067, *p* < 1 × 10^−16^) and lower overall health rating (*U* = −0.058, *p* < 4 × 10^−9^). These effects are again mediated by a DCE on BMI, which does not survive correction for multiple testing (*R* = −0.20 *±* 0.07, *U* = −0.37, *p* < 0.07), but is required by the network to explain observed effects of AFSI on the aforementioned phenotypes. While this may not represent a literal causal effect, the network structure lends insight into the results and may lend additional evidence to recent work showing that BMI-associated loci are involved in neuronal pathways linked to reward [32], or result from a latent effect of early puberty [33, 34]. It is important to note that our analysis naturally controls for confounding by socioeconomic status (SES). MR itself accounts for non-genetic confounding by SES, and bimmer does not infer that the genetic effect is either mediated by or jointly caused by “Townsend deprivation index at recruitment”. Taken together, we consider this evidence of a complex relationship between BMI and lifestyle with causal effects likely flowing in both directions.

In contrast, there are highly significant TCEs that are not reflected in the network structure. There is no path between the nodes in 2, 261 significant TCEs. Compared to connected nodes with significant TCEs, unconnected nodes tended to have fewer instruments (median instrument count 55 SNPs vs 568 SNPs for connected nodes), larger initial TCE estimates (median *R* 0.44 vs 0.07) and larger standard errors (median 0.11 vs 0.01). There are also cases where the nodes are connected, but the network estimated effect is extremely small. For example, we observe a strong TCE of past tobacco smoking on “ever taken cannabis” (*R* = −0.88 *±* 0.13, *p* < 3 × 10^−9^). Here the path between these nodes flows through BMI, followed by leukocyte count, resulting in a shrunk TCE of only −0.0002.

#### bimmer identifies concentrated sub-networks with correlated downstream effects

Our final goal was to identify densely-connected sub-networks with correlated downstream effects. To this end, we clustered the 102 phenotypes with non-zero out-degree using their outgoing shrunk TCE estimates (*U*_*i*,:_) as features. This results in clustering of phenotypes with similar downstream effects. In Figure 5, we show the genetic correlation (a), weighted-Egger TCE (b), shrunk TCE (c) and inferred DCE (d) for these 102 phenotypes as exposures and all 405 phenotypes as outcomes. This provides another view into the patterns of sharing across these matrices. This also allows us to identify several interesting sub-networks with numerous downstream effects. We selected four for further analysis, corresponding to traits associated with 1) morphology (Figure 6a), 2) blood-biomarkers (Figure 6b), 3) red blood cells (erythrocyte) (Figure 6c), and 4) heart-disease (Figure 6d). In all cases, every node within the sub-network is reachable from every other node. These sub-networks also tend to include traits that are related by definition. In many of these cases, bimmer puts a bi-directed edge between the two nodes, for example between BMI and weight. bimmer also puts a bi-directed edge between between mean sphered cell volume and mean reticulyte volume, as well as between sitting height and “predicted forced expiratory volume in one second”. While this does not happen universally, bimmer generally succeeds at identifying groups of traits which could be analyzed jointly.

**Figure 5:**
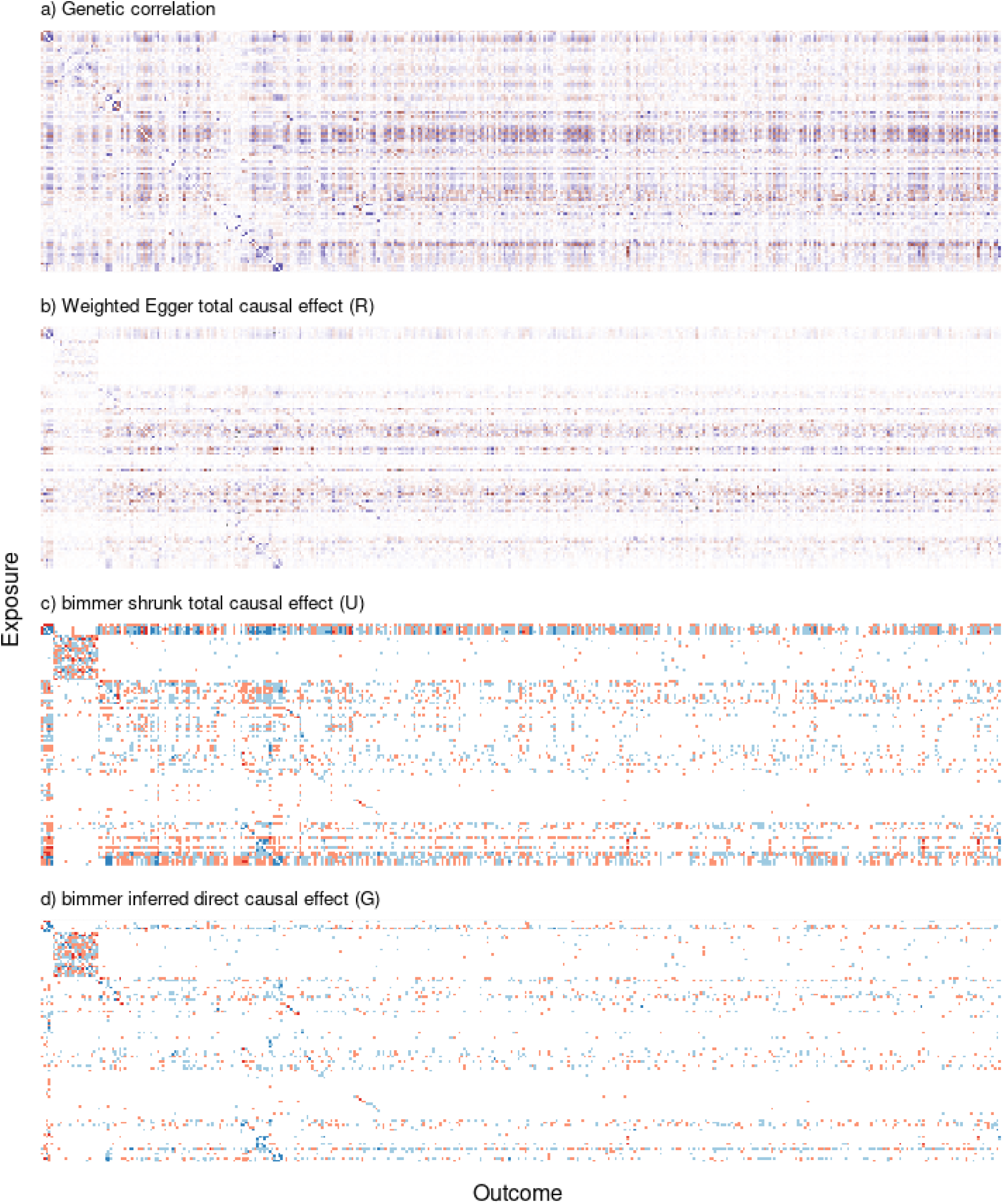
Clustering by bimmer shrunk TCE reveals groups of nodes with correlated downstream effects in the UK Biobank. We show (a) the genetic correlation, (b) the weighted Egger TCE, (c) the bimmer shrunk TCE and (d) the bimmer inferred DCE for the 102 phenotypes with non-zero out-degree (y-axis) against all 405 phenotypes (x-axis), clustered by shrunk TCE. To emphasize the smaller effects in the latter plots, we use an alternate scale which emphasizes weak positive and negative effects (0.01 to 0.1, light blue and red, respectively) and strong positive and negative effects (0.1 to 1, dark blue and red, respectively).

**Figure 6:**
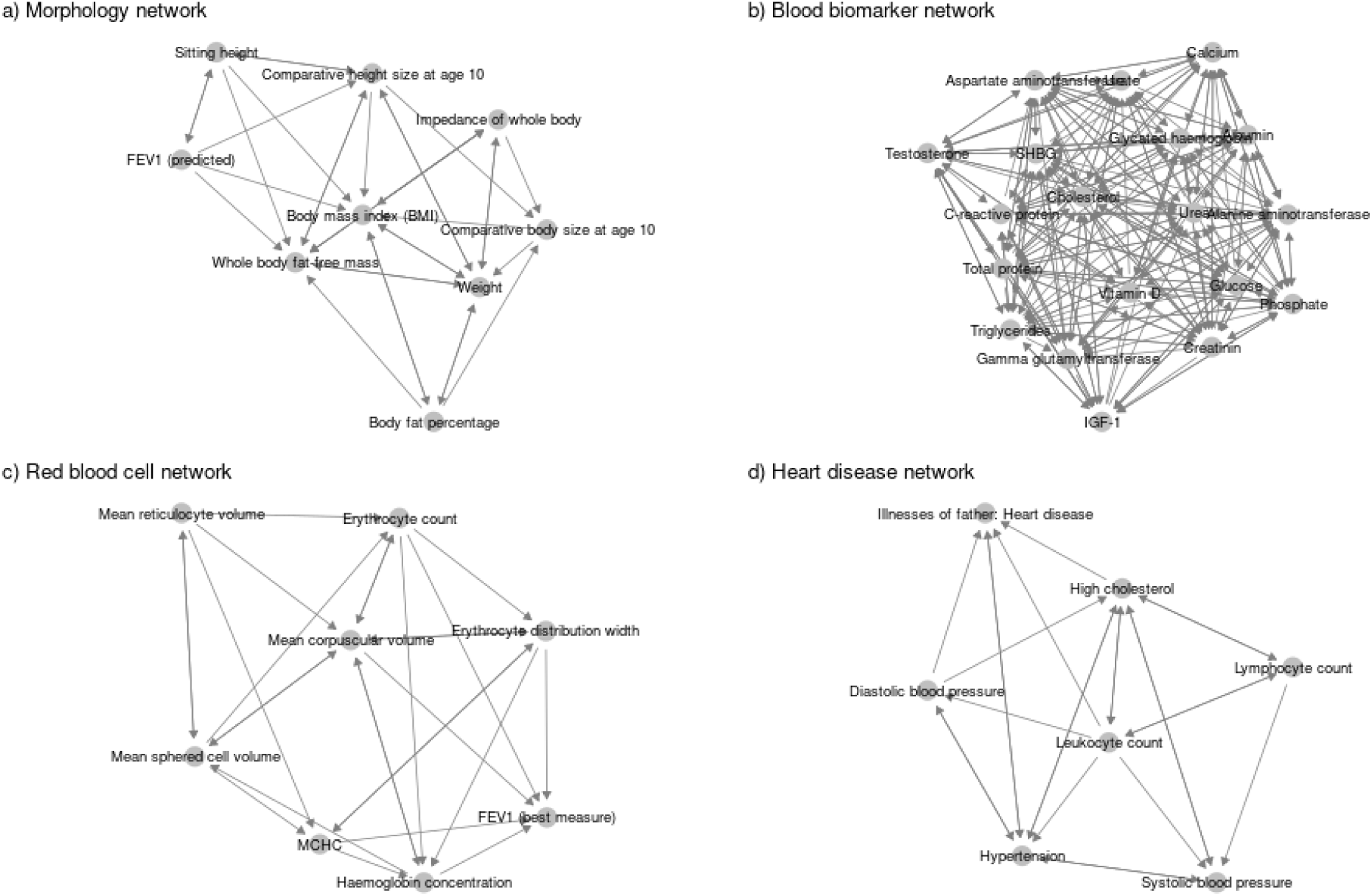
bimmer identifies densely-connected sub-networks. Clustering by bimmer shrunk TCE reveals several concentrated sub-networks, including a morphology network (a), blood biomarker network (b), red blood cell network (c) and heart disease network (d).

Part of the heart-disease network, leukocyte count had the highest overall out-degree with 244 DCEs and 177 FDR 5% significant TCEs. The heart disease network includes several well-studied phenomena including the causal effects of hypertension [35] and high cholesterol [36] on heart disease (*G* = 0.023, *p* < 3 × 10^−5^ and *G* = 0.178, *p* < 1 × 10^−16^ respectively). There is evidence for both a DCE of leukocyte count on heart disease (*G* = 0.023, *p* < 2 × 10^−3^, [37]) and indirect effects via high cholesterol (*G* = 0.051, *p* < 4 × 10^−7^) and diastolic blood pressure (*G* = 0.074, *p* < 7 × 10^−4^), both of which are previously studied pathways [38]. Interestingly, the network has an edge from leukocyte count to both systolic blood pressure and hypertension, but these do not survive correction for multiple testing as TCEs (*p* = 0.055 and *p* = 0.063, respectively). We consider this evidence of a complex mechanism by which white blood cell traits effect heart disease risk via multiple causal pathways, warranting further study. One interesting downstream effect of the heart-disease sub-network is related to choice of pain medication. We detect a positive DCE of high cholesterol on aspirin use (*G* = 0.065, *p* < 3 × 10^−6^) and a negative effect on ibuprofen use (*G* = −0.025, *p* < 0.001). This could reflect common medical advice for patients at risk of heart disease to choose aspirin, which has long been thought to reduce risk [39], and avoid ibuprofen, which is thought to reduce the effectiveness of aspirin [40]. Another interesting set of traits downstream of this sub-network are related to personality. We find evidence for a causal effect of leukocyte count on “suffer from nerves” (*G* = 0.036, *p* < 1 × 10^−10^), “worrier / anxious feelings” (*G* = 0.036, *p* < 2 × 10^−10^), neuroticism score (*G* = 0.042, *p* < 1 × 10^−7^) and “tense / highly strung” (*G* = 0.035, *p* < 5 × 10^−10^). This adds to a growing body of literature on the relationship between inflammatory biomarkers and personality [41, 42].

The blood biomarker sub-network is particularly dense, consisting of 255 direct connections, 236 of which represent significant TCEs at FDR 5%. The network implies that higher testosterone levels have numerous health consequences, many of which are related to lung function. For example, higher testosterone protects against shortness of breath (*G* = −0.028, *p* < 4 × 10^−3^) while directly increasing risk of lung cancer (i.e. not mediated through smoking, *G* = 0.013, *p* < 0.02). There is also a direct protective effect of testosterone on asthma (*G* = −0.008, *p* < 0.05) that does not survive our pruning procedure. This lends causal support to recent observational studies linking increased testosterone to lung cancer risk after controlling for smoking status [43] and mouse studies linking decreased testosterone to asthma risk [44]. We also observe an effect of testosterone on loud music exposure frequency (*G* = 0.018, *p* < 0.001). bimmer also infers a DCE of lower urea on “worrier / anxious feelings” that does not survive correction for multiple testing as a TCE (*G* = 0.011, *p* = 0.062). This lends additional evidence urea levels can have psychological consequences [45, 46, 47].

BMI has the second-highest out-degree of any phenotype considered with 215 direct effects, and 175 FDR 5% significant TCEs. Many of the strongest downstream effects of BMI are dietary in nature, including vegetable intake (*G* = 0.09, *p* < 1 × 10^−16^), milk-type (*G* = 0.15, *p* < 1 × 10^−16^), and dietary variation (*G* = 0.115, *p* < 1 × 10^−16^). These findings lend additional support to the recent literature on BMI as the cause of traits thought to lead to higher BMI (e.g. exercise [15]) and the observation that BMI-increasing genetic variants tend to be linked to genes with a role in brain function [48, 49, 32]. We consider this further evidence that causal effects between BMI and lifestyle flow in both directions. Morphology-related traits are also linked to numerous diseases, perhaps best exemplified by the strong causal effect of BMI on lower overall health rating (*G* = 0.100, *p* < 1 × 10^−16^).

In the red blood cell network, erythrocyte count and haemoglobin concentration (HC) both have high out-degree, with 117 and 132 direct effects, respectively. Erythrocyte count has numerous health consequences, for example a direct effect on lower overall health rating (*G* = 0.03, *p* < 3 × 10^−10^). Many of the top direct effects of HC involve platelet structure, for example volume of blood occupied by platelets (plateletcrit, *G* = −0.13, *p* < 3 × 10^−11^) and platelet count (*G* = −0.15, *p* < 2 × 10^−10^). Interestingly, our model predicts a direct effect of HC on bleeding gums (*G* = 0.036, *p* < 2 × 10^−9^); that is, one that is not mediated by the aforementioned effects on platelets. This may reflect a lack of power to detect the direct effect of platelets on bleeding gums, or the existence of an alternative pathway. Finally, we detect a DCE of HC on cardiac arrythmia (*G* = −0.042, *p* < 5*e* 9), lending causal support to a recent population-based study linking HC and atrial fibrillation [50].

## 3 Discussion

As biobanks continue to grow in size and scope, new methods that are able to leverage their power while overcoming common pitfalls are required. These datasets offer unprecedented opportunity to study the causal relationship between biomarkers, complex traits and diseases. Here, we have introduced bi-directional mediated Mendelian randomization (bimmer), a novel approach to inferring sparse networks of direct causal effects from phenome-scale GWAS summary statistics. We have shown through extensive simulations that bimmer is able to learn many kinds of causal graph structures even in the presence of non-causal genetic correlation and differential power across phenotypes. We have demonstrated that our method enables analyses that would otherwise be impossible. For example, we are able to interrogate the complexity of the network by analyzing the path length distribution and proportion of effect explained by the shortest path. We are also able to identify densely connected sub-networks with correlated downstream effects. By applying our method to the UK Biobank, we lend causal support to several recent observational studies.

Generally speaking, causal claims should be backed by thorough analysis resulting from multiple studies with differing assumptions and input from domain experts. This raises the question of whether phenome-scale causal inference, where the number of pairs of traits to be tested renders this unrealistic, is even possible. Instead, in this setting one should focus on causal network discovery, learning putatively causal structures that can suggest avenues for further work. We have demonstrated that causal discovery is invaluable for understanding the results of phenome-wide Mendelian randomization. Simultaneous inference of edges in the causal graph can prioritize seemingly-insignificant connections that are globally relevant, remove false-positives with significant TCE *p*-values that cannot be coherently incorporated into the network, and enable interpretation of surprising TCEs in the context of the network structure.

Our approach is conceptually simple and can be viewed both as a method and a framework. First, we calculate a TCE matrix. This is analogous to a correlation matrix, except that it is not symmetric and its entries represent apparent causal effects rather than correlations. Then, we find a sparse approximate inverse to this matrix, which represents a causal graph. This is analogous to glasso, except that we produce a directed graph instead of an undirected one. This means that bimmer can use any MR method that is able to produce bi-directed effect estimates, allowing researchers to choose the method that best accommodates the assumptions of the setting they work in. It also allows bimmer to naturally accommodate other potential approaches to convert the TCE matrix into a causal network. While we are not aware of other methods for this specific problem, our method contributes to the extensive fields of biological and causal network discovery. For example, Frot *et al.* [8] combines a low-rank plus sparse decomposition of the covariance matrix [9, 7] with the algorithm *intervention calculus when the DAG is absent* [51] to learn directed acyclic causal graphs. There, the low-rank component is interpreted as a set of latent variables capturing unobserved confounding, whereas our method uses Mendelian randomization for this task. Because of the modular nature of our approach, inspre could be easily extended to include a low-rank component which could help capture residual confounding. Our method is also related to *silencing* [52]. There, the authors use physical arguments pertaining to perturbations at equilibrium to derive a formula similar to (1). However, they use an exact inversion which leads to poor performance in practice [53]. Finally, Dahl *et al.* [54] and Stegle *et al.* [10] model a low rank component in the observed data to capture population structure and latent gene expression factors, respectively. These methods require access to the primary data, learn undirected graphs, and do not scale to biobanks.

Our approach also builds on recent MR literature. In particular, multi-variable Mendelian randomization methods are able to compute direct causal effects when there are multiple potential exposures and a single outcome [29, 28]. While these methods work well in that setting, we have shown that they are not well-suited to the more general network inference problem that we consider here. Another approach, network Mendelian randomization, calculates the effect of an exposure on an outcome while accounting for the effects of a third variable [17]. Our method can be thought of as a generalization of this approach to an arbitrary number of phenotypes without pre-specifying any as exposures or outcomes. To calculate the TCE matrix, we use a novel approach to bi-directional MR with Egger regression weights that reduce the effect of pleiotropic SNPs. This is related to several recent methods. In particular, gwas-pc uses asymmetry in the effect size distributions to choose an effect direction between the two phenotypes. Similarly, LCV uses this asymmetry to fit a latent variable model, where imbalanced genetic correlation between the phenotypes and latent variable imply the effect direction. Compared to these methods, our approach offers several advantages. First, like LCV but unlike gwas-pc, our method controls the type-I error rate when there is non-causal genetic correlation and differential power. Second, like gwas-pc, but unlike LCV, our method estimates a quantity that is interpretable as the effect of one phenotype on the other. Finally unlike both, we are able to estimate both effect directions simultaneously, allowing our model to accommodate graphs with cycles. Our method is also related to the recently proposed CAUSE [55]. There, the authors model observed SNPs effects as coming from a mixture of pleiotropic and non-pleiotropic SNPs. Our method can be viewed as a simple and fast approximation to this method where SNPs that appear pleiotropic are given a lower regression weight.

However, our approach does have some weaknesses. First, our method requires that we split the initial cohort into instrument discovery and effect estimation sub-cohorts. This is common in MR methods, but LCV has the distinct advantage of using all SNPs, which obviates the need for sample splitting and should improve power. Second, while there are some phenotype pairs where a direct cause makes sense, there are others where causality is almost certainly better interpreted as the action of a latent variable. Indeed, it is likely that some of the causal effects we infer actually represent shared causal pathways. Finally, our method suffers from a modest increase in false positives in the *B* → *A* direction when there is a strong effect from *A* → *B*. When there are many causal SNPs and the effect of *A* on *B* is strong, some SNPs that directly effect *A* can be mistakenly used as instruments for the effect of *B* on *A*. While our method reduces the magnitude of this estimated effect, it can still give some false positives. Our approach also still suffers from weak instrument bias, generally underestimating the causal effect, which reduces power [56].

The second step of our method involves finding a sparse inverse to a noisily measured matrix, and is therefore closely related to the graphical lasso. Like glasso, our method has a single regularization parameter that can be set in a straightforward manner. However, a key advantage of our approach is that we are able to incorporate observation weights. This is extremely important in our application since the standard errors of the TCE matrix can vary dramatically. This also allows us to approximately invert matrices with missing data, implicitly performing matrix completion by leveraging assumed sparsity in the inverse. This enables stability-based selection of lasso parameter *λ* without access to the underlying data by using random masks. There are other approaches to weighted graphical lasso [57, 58], however these weights represent prior knowledge and not statistical uncertainty. Here again, our method could easily accommodate prior biological knowledge through a simple modification of the LASSO penalty. We found that for many classes of graphs, inspre and glasso produced similar results, however there were some settings where glasso clearly performed better and vice versa. Moreover, our method is substantially slower than glasso. If *κ* is the number of iterations required to reach convergence, glasso very roughly requires time *O*(*κD*^3^), whereas inspre requires time roughly *O*(*κD*^4^) for *D* phenotypes. We suspect that there may be ways of improving the speed of our approach, and in spite of these limitations, the novel capabilities of inspre suggest it may find utility outside the scope of MR.

Importantly, bimmer only requires GWAS summary statistics. This allows us to apply our method to the realistic setting where each phenotype is only measured on a subset of individuals. While the UK BioBank primary genotypes and phenotypes are readily available, this is not the case for many cohorts. Summary statistics are both legally and practically easier to share, and faster to work with when the primary data is large [59]. They also enable researchers to work with data from a standardized analysis pipeline, such as [30]. Strictly speaking, our method does not even require summary statistics. If MR analysis results are already available for every pair of a set of phenotypes, one can use them to construct the matrix *R* and then infer *G* with inspre. In this setting, it is of paramount importance that the researcher verify the underlying studies were conducted in a way to minimize the effect of horizontal pleiotropy.

In this work we have begun to elucidate the connection between Mendelian randomization and the omnigenic model [1]. The effects of genetic variants can be used to find and orient edges in the DCE graph underlying trait variation, and long-range effects can be modeled as paths in this sparse graph resulting in ubiquitous but small effects. Our method can be applied well beyond the scope considered here. We are particularly interested in the application to datasets of molecular phenotypes. These datasets generally have much smaller sample sizes, but molecular phenotypes also tend to have larger, localized SNP effect sizes [60], which improves the efficiency of MR. Inverse sparse regression could also be applied to datasets from high-throughput CRISPR-based genetic perturbation experiments to separate out direct effects from mediated regulatory relationships. We look forward to pursuing these avenues in future work.

## 4 Methods

### 4.1 Trait model

Our goal is to estimate a sparse graph of direct causal effects (DCE), *G*, from summary association statistics between genotypes *X* and phenotypes *Y*. We model the SNP effects *β* and the causal graph as fixed effects, and assume that the genotypes *X* are sampled independently from a population. For convenience, we assume that SNPs and phenotypes have been normalized to have mean 0 and variance 1. We also assume that SNPs are uncorrelated (no linkage disequilibrium, LD) and use LD-pruned variants in all analyses of real data. Our model is

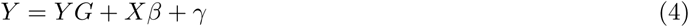

where *γ* represents unmeasured factors. Let *R* be the *D* × *D* matrix of estimated TCEs from MR. Our goal is to show that under this model *G* = *I* − *R*^−1^*D*[1*/R*^−1^]. If *I* − *G* is invertible, (4) can be re-written

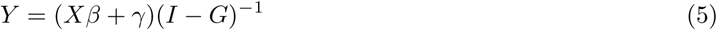

For the purposes of this derivation, we consider solving (5) using two-stage least-squares with *X* as instruments, but in practice any MR method can be used. For now, we assume each SNP acts only on one phenotype (there is no pleiotropy) and that we know which phenotype it is. First we regress each instrument on its phenotype and use these effect estimates to calculate a set of phenotype scores for each individual. Next, we regress each phenotype score on the observed values of the other phenotypes, creating a matrix containing estimates of the total causal effect (TCE) of each phenotype on every other. This gives the estimated effect matrix 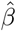. Using (5), we can find 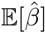,

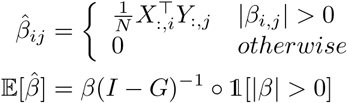

where 𝟙 is an indicator function and ∘ is the Hadamard matrix product. Using (4), the TCE matrix is

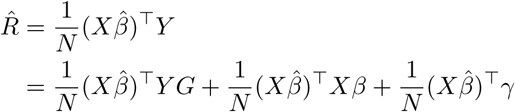

Taking expectations and assuming that the environment random effect *γ* has 0 mean we obtain,

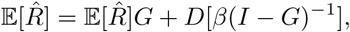

where the diagonal operator 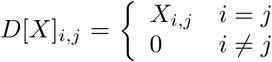 sets off-diagonal elements of a matrix to 0. Since 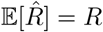, this tells us that *R* satisfies the recurrence *R* = *RG* off the diagonal, from which it follows that [53],

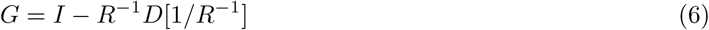

where */* indicates elementwise division.

In practice we don’t know which SNP effects which phenotype, and there can be correlated pleiotropic effects. Consider a pair of phenotypes *i* and *j* where phenotype *i* has a direct causal effect of *G*_*i,j*_ on phenotype *j*. If SNP *k* has a direct effect on phenotype *i* of size *β*_*k,i*_, but no direct effect on phenotype *j* (i.e. no pleiotropy) then the observed effect of SNP *k* on phenotype *j* is

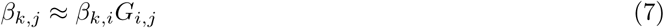

This SNP therefore contributes 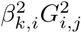 to the variance of *Y*_*j*_, whereas pleiotropic SNPs will contribute 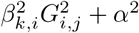 for some pleiotropic effect size *α*. Therefore, SNPs that appear to have a larger absolute effect on the exposure relative to the outcome in a discovery cohort are more likely to satisfy (7). First, we split the samples into two sets and generate two sets of summary statistics, one for SNP discovery and weight estimation (the discovery set) and one for TCE estimation (the estimation set). Using the discovery set, we identify the set of SNPs marginally associated at p-value threshold *p* for each phenotype *i*. Call this set 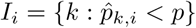. For every SNP *k* ∈ *I*_*i*_ and every phenotype *j*, we calculate the Welch test statistic for a two sample difference in mean with unequal variances [24]

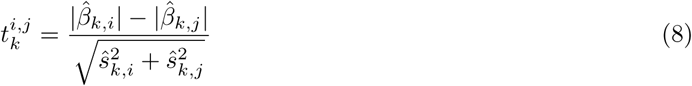

and use this to construct a weight 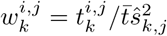.

### 4.2 Inverse sparse regression

If we knew *R* exactly, we could simply invert it and plug the inverse into (6). However, we only have access to the noisy estimate 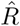, which is not necessarily well-conditioned or even invertible. Instead, we assume that the underlying directed graph of DCE is sparse. We observe that in (6), *G* is sparse if and only if 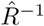 is sparse, and so we can think of solving (6) as finding a sparse matrix inverse. Let *A* be an arbitrary *D* × *D* matrix (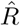 in bimmer). We seek matrices *U, V* with *V U* = *I* that minimize the loss,

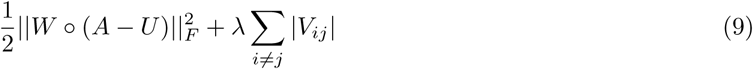

We minimize this loss using alternating direction method of multipliers (ADMM) [61]. Let Θ^*k*^ be a matrix of Lagrange multipliers. The updates for *U* ^*k*^, *V* ^*k*^ and Θ^*k*^ are

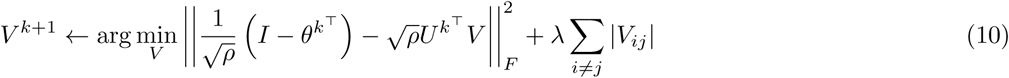

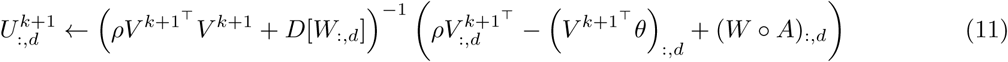

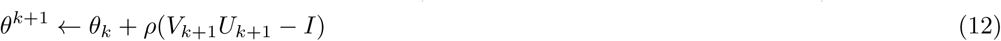

where *ρ* is the penalty parameter [61]. The update for *V* ^*k*+1^ is a straightforward LASSO regression. For the update for *U* we use the biconjugate gradient stabilized method implemented in the Rlinsolve package to solve the linear system rather than explicitly computing the inverse [62]. We always start from the initial condition *U*_0_ = *V*_0_ = *I*. For the derivation of these equations including the specifics of how we tune the penalty parameter see the Supplemental note.

### 4.3 Setting the LASSO penalty using stability selection

We use an adaptation of the Stability Approach to Regularization Selection (StARS, [23]) to select the regularization parameter. StARS leverages the intuition that smaller values of *λ* yield graphs that are more stable under random re-samplings of the input data to construct an interperatable quantity representing the average probability that each edge is included in the graph for each value of *λ* in (9). Let *ϕ*_*λ*_ be a *D* × *D* matrix where entry *i, j* is the probability that each edge *i, j* is included in the graph for regularization parameter *λ*. Our goal is to estimate *ϕ*_*λ*_ for many choices of *λ* and turn this into a graph instability measure *D*_*λ*_. Let 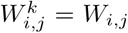 with probability *p* and 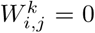 with probability 1 − *p*. Let 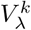 be the approximate inverse of *A* resulting from fitting (9) for regularization setting *λ* and weight set *W* ^*k*^. Let 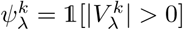. Then *ϕ*_*λ*_ can be estimated as

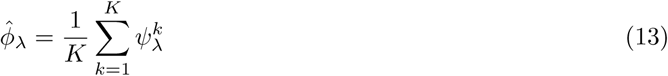

using *K* independent random masks. The instability measure *D*_*λ*_ is estimated as [23]

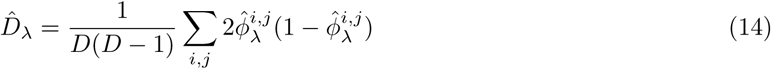

Clearly, *D* = 0 for very large values of *λ*, where 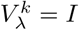 for every mask *k*. As *λ* becomes smaller, *D* rises, but as *λ* approaches 0, *D* → 0 as 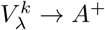. Following [23], we first normalize 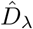 by setting it to 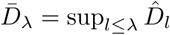 and then choose the smallest value of *λ* with stability below a cut point 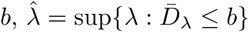.

## 5 Code Availability

All code used in the production of this manuscript is available at https://github.com/brielin/bimmer and https://github.com/brielin/inspre. The full data analysis results are available at https://zenodo.org/record/3895125.

## 6 Acknowledgments

BCB would like to thank Harold Pimmentel and Andy Dahl for helpful feedback on the manuscript. BCB would also like to thank Lior Pachter and Nicolas Bray for insightful discussion of the proposed method. DAK would like to thank Jonathan Pritchard for useful feedback on the initial concept. BCB is funded by post-doctoral fellowship from the Data Science Institute at Columbia University.

## 7 Author Contributions

BCB and DAK jointly formulated the model and estimation procedure. BCB wrote the code, conducted analyses, and drafted the manuscript. DAK supervised and assisted with editing the manuscript.

## Supplemental Note

### Alternating direction method of multipliers

First, consider the unweighted optimization problem

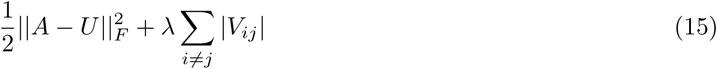

The augmented Lagrangian is,

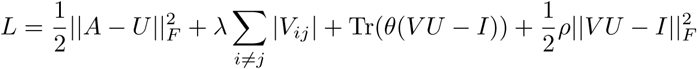

The update for *V* can be found by noticing that minimizing *L* is equivalent to solving a lasso regression with design matrix 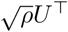 and response 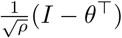,

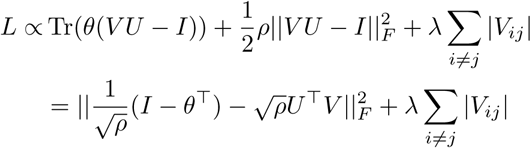

The update for *U* can be found by taking the gradient ∇_*U*_ *L* and setting it to 0,

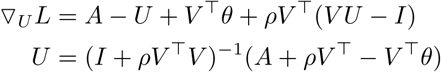

ADMM gives the update for *θ* [61],

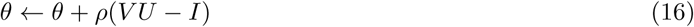

Now we consider the weighted version. Assume that in addition to the matrix *A*, we also have a matrix of standard errors of the entries of *A, S*_*A*_. Let 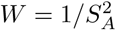 be a matrix of inverse variance weights. We now seek matrices *U, V* with *V U* = *I* that minimize the loss,

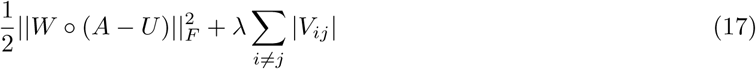

This does not effect the update for *V*, however the gradient of the augmented Lagrangian with respect to *U* is now,

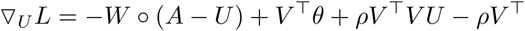

which separates over columns of *U*, giving the update

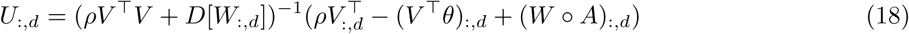

where here the *D* operator creates a matrix with *W*_:,*d*_ on the diagonal and 0 elsewhere.

ADMM also requires that we set the parameter *ρ*, which controls the balance in the objective between the primal and dual constraints [61]. We follow standard practice of setting rho to an initial value and increasing or decreasing it according to the ratio of the solution to the primal and dual feasibility constraints. The primal residual at iteration *k* + 1 is given by *r*^*k*+1^ = *V* ^*k*+1^*U* ^*k*+1^ −*I*. The dual residual is found by setting ∇_*U*_ *L*^*k*^ = 0 and evaluating it at *U*_*k*+1_

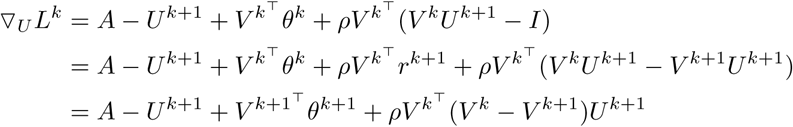

Therefore the dual residual is [61]

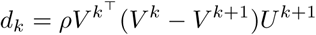

and we can adjust *ρ* as follows,

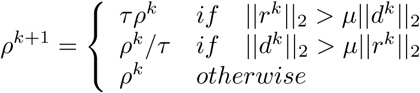

which reduces the impact of the initial choice of *ρ*. While this may appear to be a lot of parameters, they effect the convergence of the algorithm substantially more than the solution obtained. We always use the default values *ρ* = 10, *µ* = 10, *τ* = 2.

---

**Figure S1:**
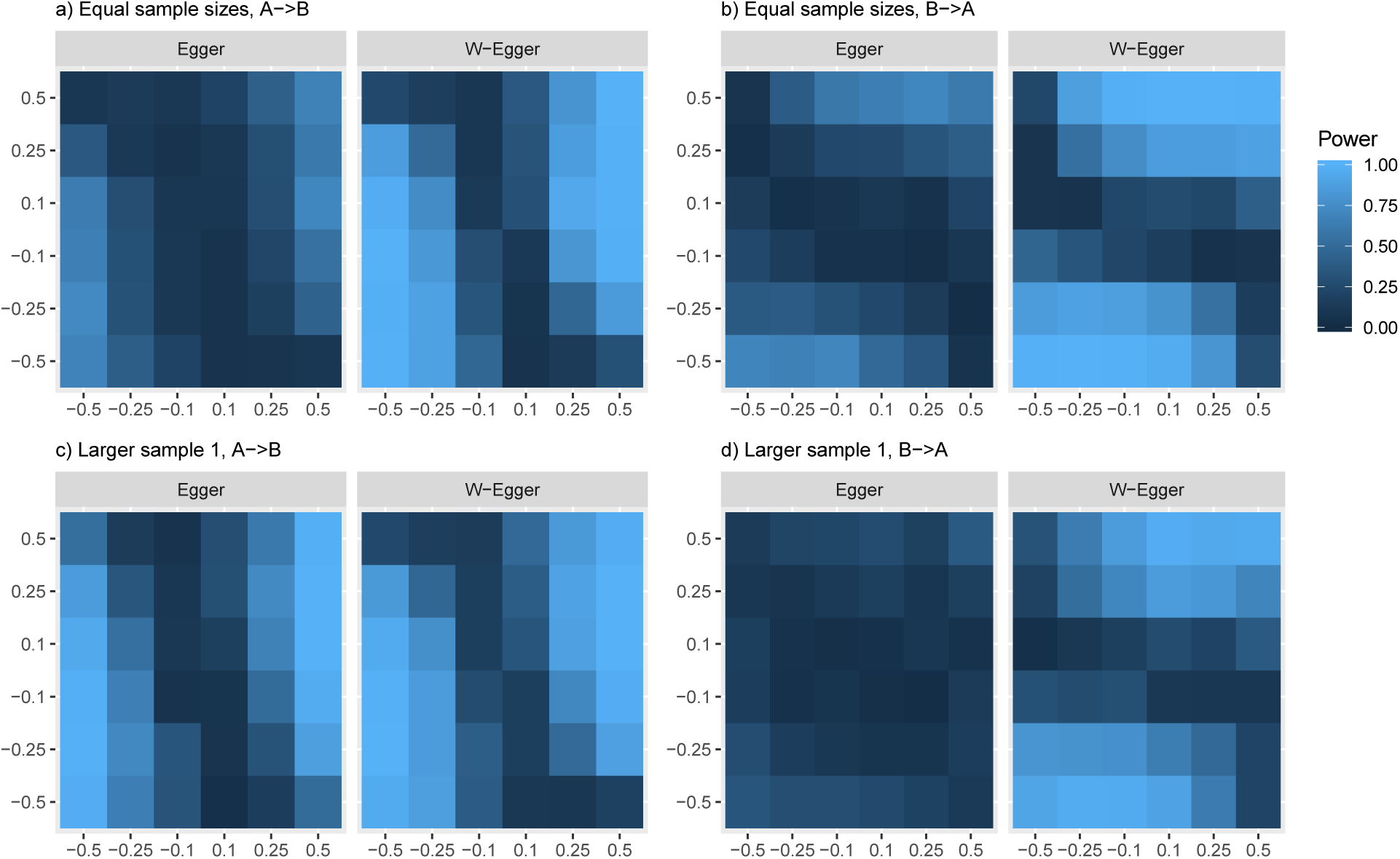
Weighted Egger regression improves power under the two-way alt. We simulated GWAS summary statistics for two phenotypes (*A, B*) with *M* = 1, 000, 000 independent SNPs, 20% heritability and *N* = 100, 000 individuals in both the SNP discovery and effect estimation cohorts. In each simulation, there were 5, 000 causal SNPs per phenotype of which 1, 000 were shared with uncorrelated effect sizes. a) Power to detect the effect of *A* on *B* when the studies have equal sample sizes. Our weighting scheme increases power of standard Egger regression, but both methods struggle to detect when the traits cancel each other out. b) Power to detect the effect of *B* on *A*. Our approach improves power and the cancellation pattern is transposed. c) Power to detect the effect of *A* on *B* when *A* has a larger sample size. Our approach improves power, though both do well. d) Power to detect the effect of *B* on *A* when *A* has a larger sample size. Our approach improves power substantially over standard Egger regression, which struggles to detect the effect. Results are the average of 250 simulations per pair of effects.

**Figure S2:**
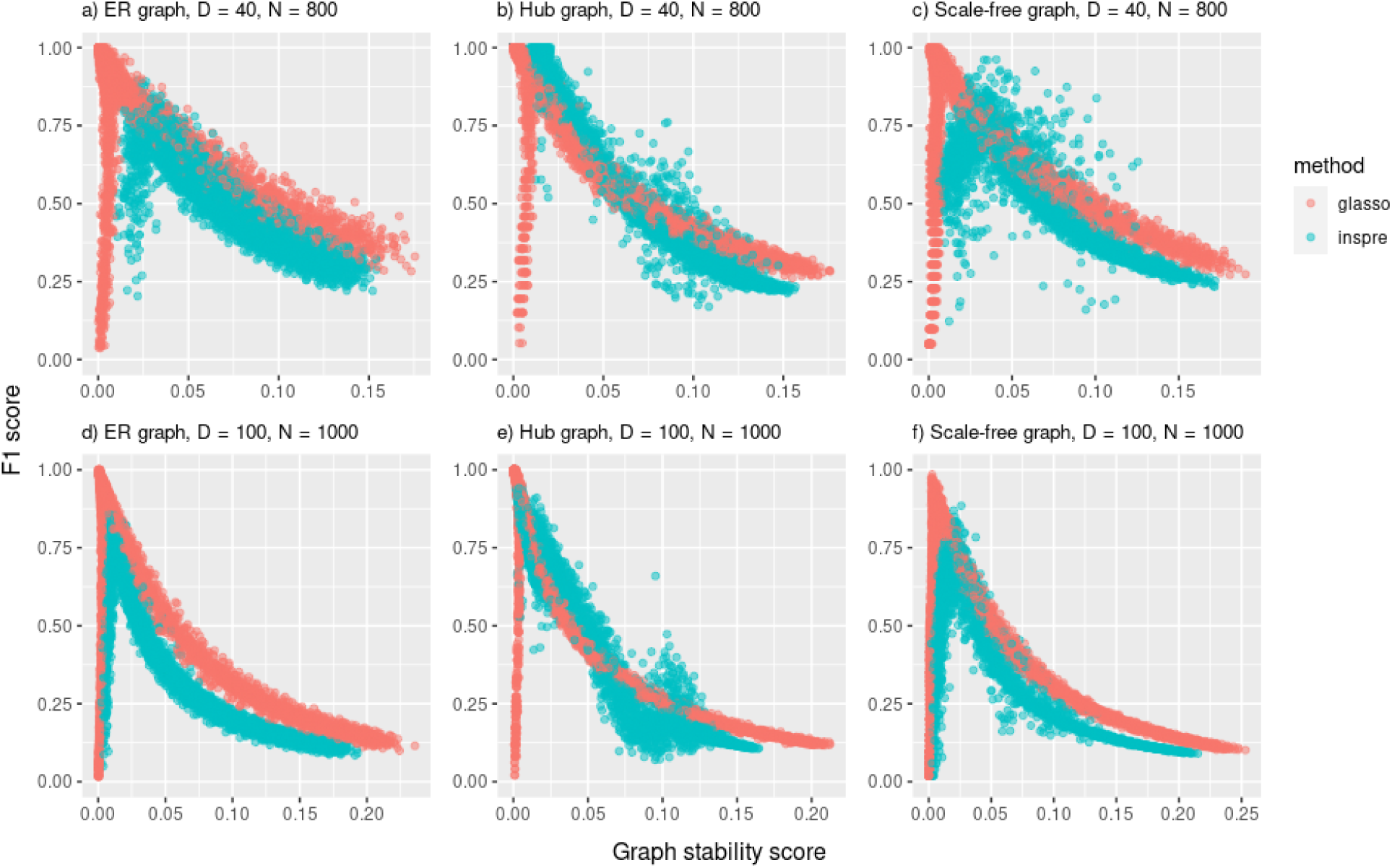
inspre performs similarly to glasso on data from gaussian graphical models. We simulated data from a multivariate normal distribution with a sparse precision matrix for various sample sizes, dimensionalities and graph structures. Then we evaluated the relationship between *F*_1_-score and graph stability for both inspre and glasso. a) inspre and glasso perform similarly for Erdos-Reyni graphs, b) hub graphs and c) scale-free graphs with 40 dimensions and 800 samples. d) At 100 features and 500 samples, glasso outperforms inspre on Erdos-Reyni graphs, but the opposite is true for hub graphs (e). f) glasso also slightly outperforms inspre on scale-free graphs in this setting.

**Figure S3:**
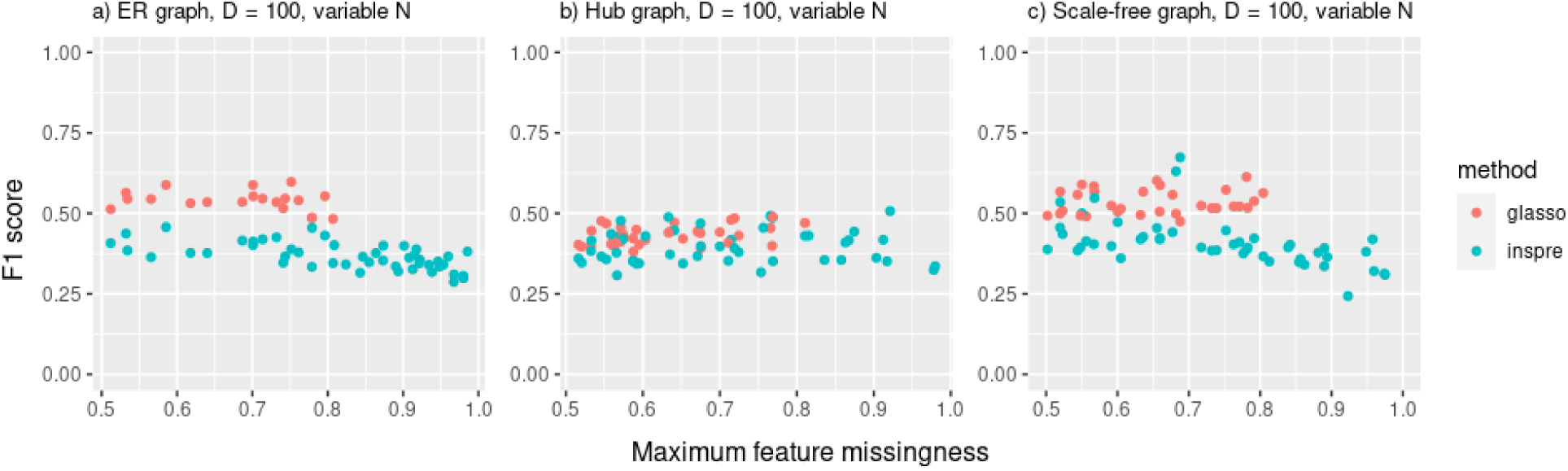
inspre is able to produce results when there is differential sample size across features while glasso diverges. We simulated data from a gaussian graphical model with 100 features and samples sizes ranging from 20 −2000 per feature. inspre continues to produce results when some features have only 20 −400 samples, while glasso needs at least 400 for a) Erdos-Reyni, b) hub and c) scale-free graphs.

**Figure S4:**
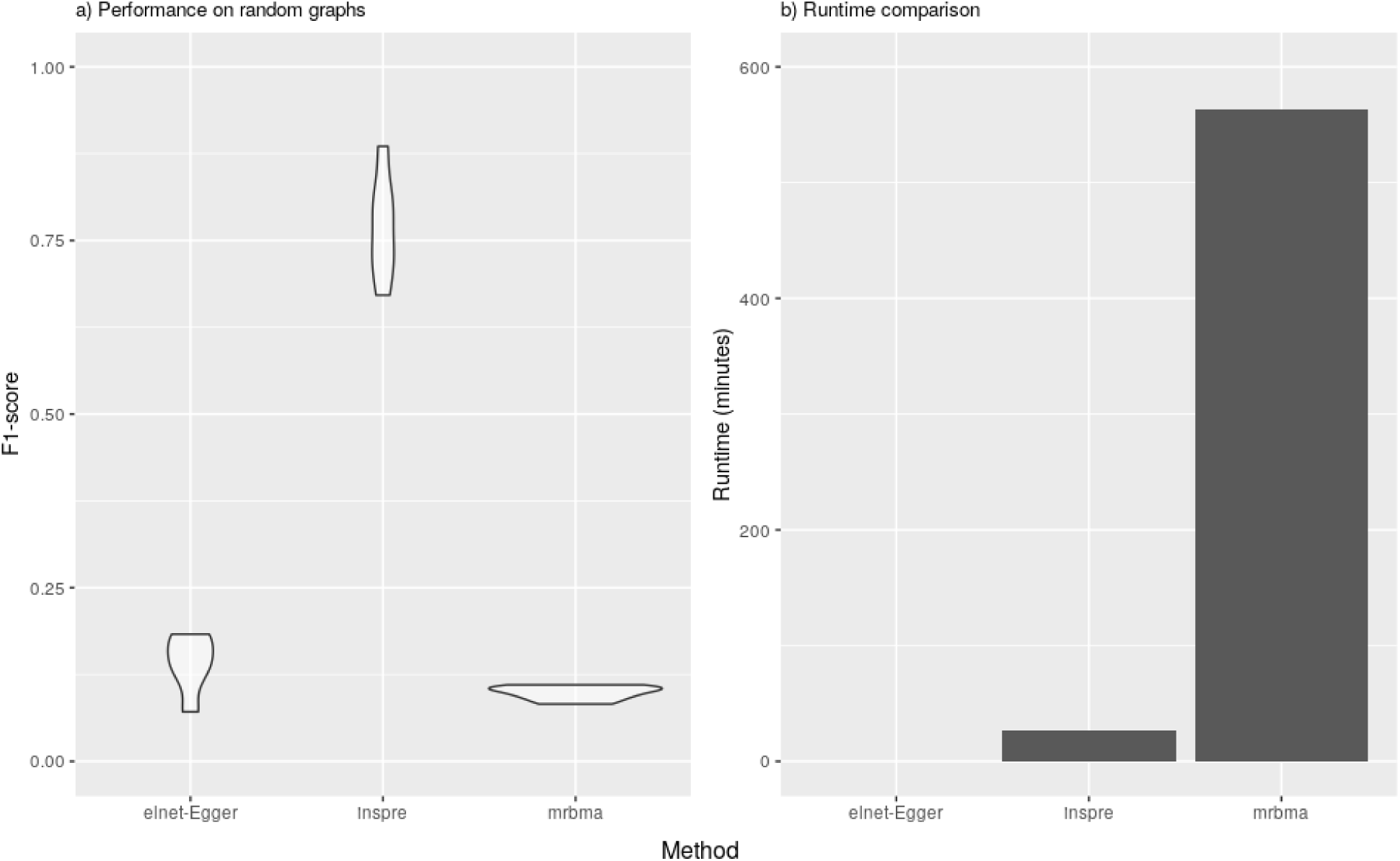
bimmer accurately infers small random graphs much faster than MR-BMA. We simulated Summary statistics for 40 phenotypes with 3000 causal effects each, 1000 of which were shared with uncorrelated effects per pair of phenotypes. a) inspre outperforms both elnet-Egger and MR-BMA, which takes about 20 times longer to run than inspre (b).

**Figure S5:**
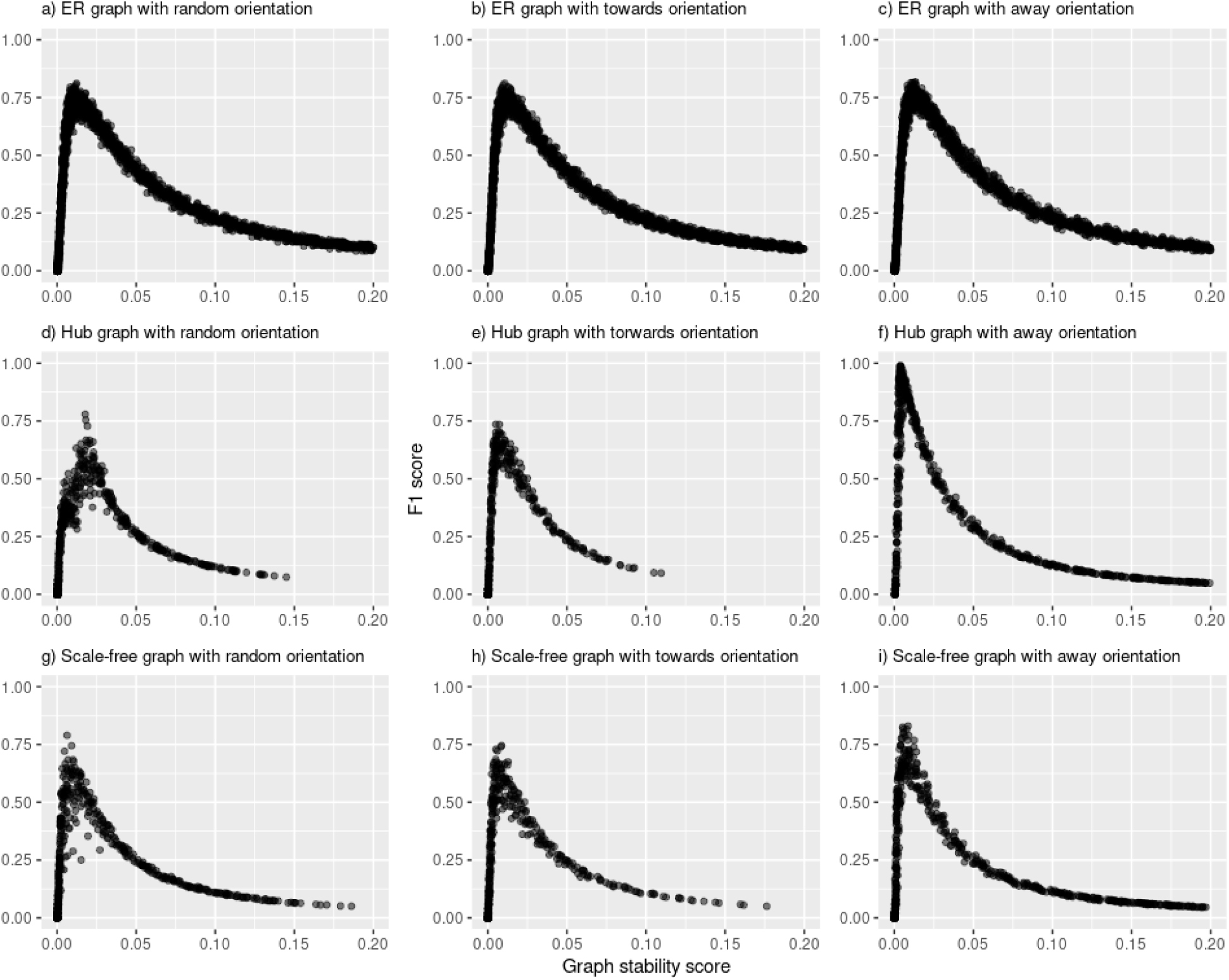
bimmer accurately infers the causal graph for many graph structures and node orientations. We simulated summary statistics for 100 phenotypes with 3000 causal effects each, 1000 of which were shared with uncorrelated effects per pair of phenotypes. We varied the structure and edge orientation of the causal graph underlying the phenotypes. We show the *F*_1_-score of the method against the stability score for a) Erdos Reyni graphs with randomly oriented edges, b) Erdos-Reyni graphs with edges oriented towards high-degree nodes, c) Erdos-Reyni graphs with edges oriented away from high-degree nodes, d) hub graphs with randomly oriented edges, e) hub graphs with edges oriented towards high-degree nodes, f) hub graphs with edges oriented away from high-degree nodes, g) scale-free graphs with randomly oriented edges, h) scale-free graphs with edges oriented towards high-degree nodes, i) scale-free graphs with edges oriented away from high-degree nodes.

**Figure S6:**
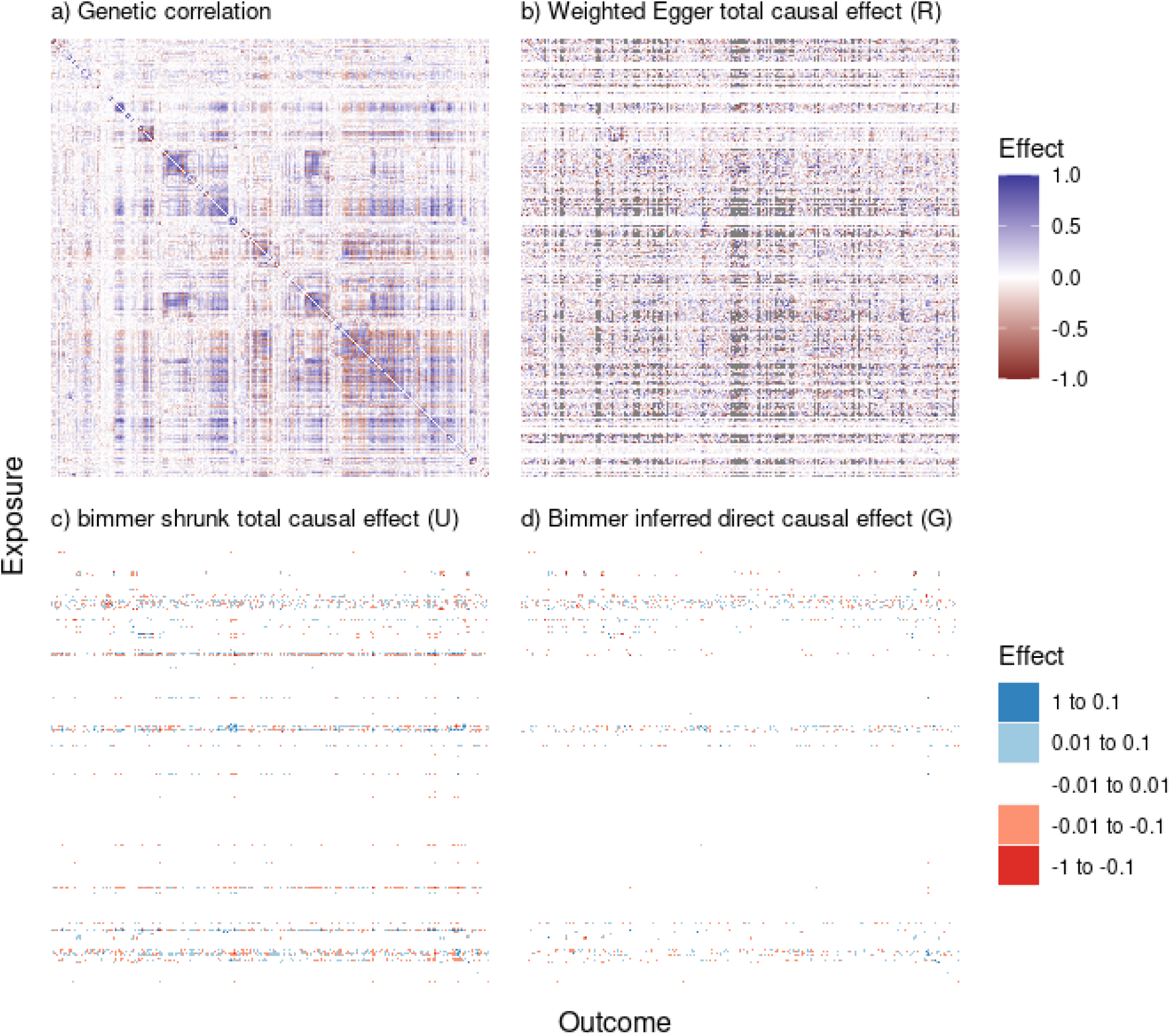
bimmer shrunk total causal effects are only weakly correlated with genetic correlation. a) Genetic correlation between all pairs of phenotypes, clustered by absolute correlation. b) Weighted Egger TCE estimates using the same clustering scheme reveals some similar patterns, but lacks the well-defined structure of the genetic correlation estimates. The bimmer shrunk TCE (c) and DCE (d) reveal a banded structure where few nodes have high out-degree with many downstream effects, but most nodes have no downstream effects (empty rows).

**Table S1:** Weighted Egger regression reduces false positives in bi-directional MR under the two way null. We simulated GWAS summary statistics for two phenotypes (*A, B*) with *M* = 1, 000, 000 independent SNPs, 20% heritability and *N* = 100, 000 individuals in both the SNP discovery and effect estimation cohorts. In each simulation, there were 5, 000 causal SNPs per phenotype and neither phenotype had an effect on the other. In the first setting pleiotropic effects are uncorrelated and all methods are well behaved. In the next setting the 1000 shared SNPs have equal effects on both phenotypes. Here Egger regression results in excess false positives which our weighting scheme reduces. In the last setting shared SNPs again have equal effects on both phenotype, but shared SNPs explain a larger proportion of the variance in the second cohort, which also has a smaller sample size. Here Egger regression results in numerous false positives, which our weighting scheme corrects. Values reflect averages over 1, 000 simulations.

| Method | $FPR_A$ | $SE_A$ | $FPR_B$ | $SE_B$ | $MAE_A$ | $SE_A$ | $MAE_B$ | $SE_B$ |
| --- | --- | --- | --- | --- | --- | --- | --- | --- |
| **Null: Uncorrelated pleiotropy** |  |  |  |  |  |  |  |  |
| Oracle | 0.041 | 0.006 | 0.054 | 0.007 | 0.009 | 0.000 | 0.009 | 0.000 |
| W-Egger | 0.053 | 0.007 | 0.058 | 0.007 | 0.042 | 0.001 | 0.042 | 0.001 |
| Egger | 0.053 | 0.007 | 0.053 | 0.007 | 0.091 | 0.002 | 0.090 | 0.002 |
| **Null: Correlated pleiotropy** |  |  |  |  |  |  |  |  |
| Oracle | 0.043 | 0.006 | 0.038 | 0.006 | 0.009 | 0.000 | 0.009 | 0.000 |
| W-Egger | 0.051 | 0.007 | 0.068 | 0.008 | 0.044 | 0.001 | 0.047 | 0.001 |
| Egger | 0.095 | 0.009 | 0.084 | 0.009 | 0.172 | 0.004 | 0.166 | 0.004 |
| **Null: Correlated pleiotropy, unequal power** |  |  |  |  |  |  |  |  |
| Oracle | 0.052 | 0.007 | 0.036 | 0.006 | 0.014 | 0.000 | 0.006 | 0.000 |
| W-Egger | 0.087 | 0.009 | 0.029 | 0.005 | 0.053 | 0.001 | 0.080 | 0.002 |
| Egger | 0.284 | 0.014 | 0.492 | 0.016 | 0.178 | 0.003 | 0.648 | 0.010 |

**Table S2:**
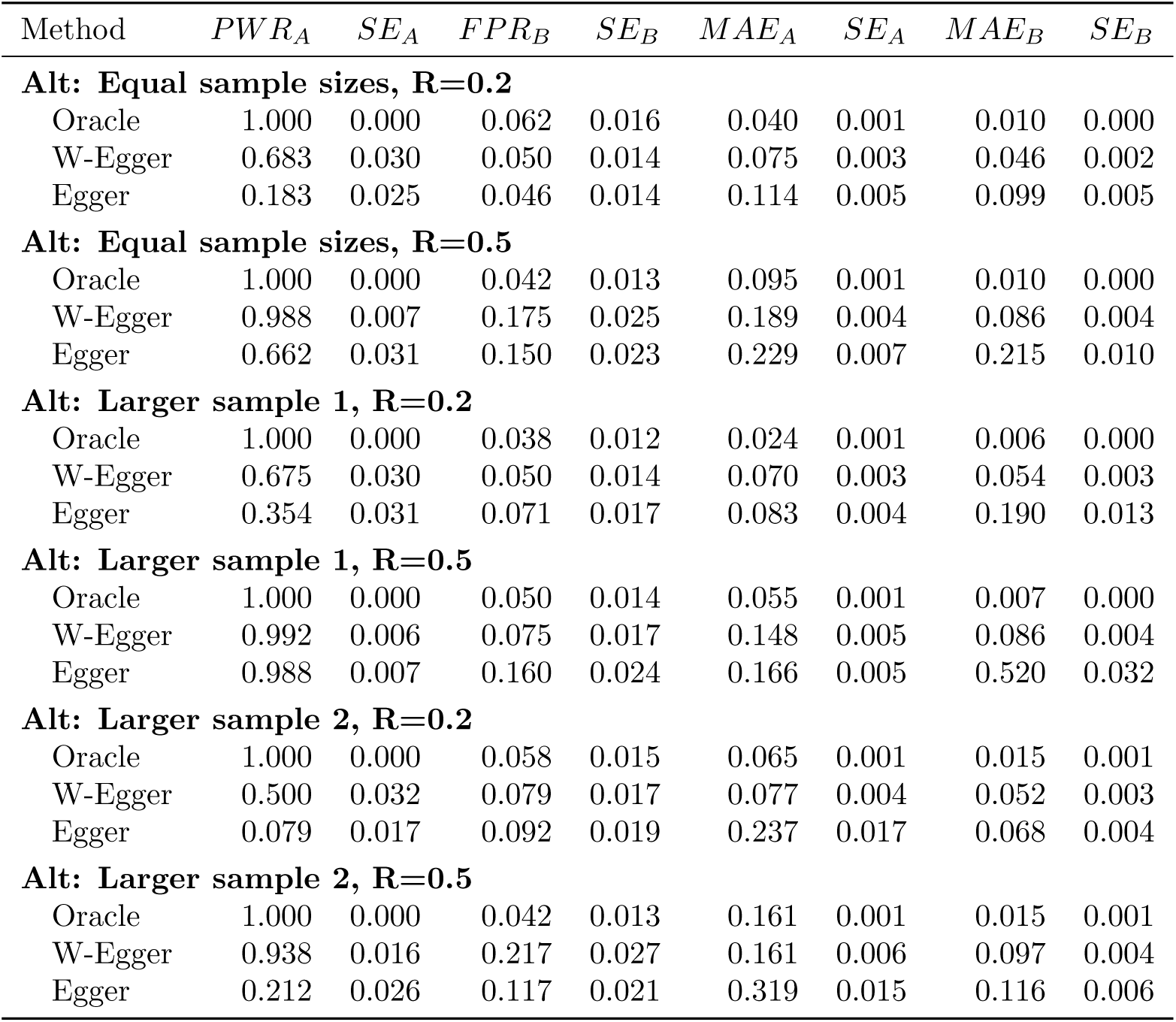
Weighted Egger regression improves power and reduces false positives under the one-way alternate hypothesis. We simulated GWAS summary statistics for two phenotypes (*A, B*) with *M* = 1, 000, 000 independent SNPs, 20% heritability and *N* = 100, 000 individuals in both the SNP discovery and effect estimation cohorts. In each simulation, there were 5, 000 causal SNPs per phenotype and *A* has a variable effect on *B*. When the cohorts have the same sample size, weighted Egger regression improves power in the alt direction while reducing the magnitude of the effect inferred in the null direction for both. This continues to hold when cohort *A* is larger and when cohort *B* is larger.

| Method | $PWR_A$ | $SE_A$ | $FPR_B$ | $SE_B$ | $MAE_A$ | $SE_A$ | $MAE_B$ | $SE_B$ |
| --- | --- | --- | --- | --- | --- | --- | --- | --- |
| **Alt: Equal sample sizes, R=0.2** |  |  |  |  |  |  |  |  |
| Oracle | 1.000 | 0.000 | 0.062 | 0.016 | 0.040 | 0.001 | 0.010 | 0.000 |
| W-Egger | 0.683 | 0.030 | 0.050 | 0.014 | 0.075 | 0.003 | 0.046 | 0.002 |
| Egger | 0.183 | 0.025 | 0.046 | 0.014 | 0.114 | 0.005 | 0.099 | 0.005 |
| **Alt: Equal sample sizes, R=0.5** |  |  |  |  |  |  |  |  |
| Oracle | 1.000 | 0.000 | 0.042 | 0.013 | 0.095 | 0.001 | 0.010 | 0.000 |
| W-Egger | 0.988 | 0.007 | 0.175 | 0.025 | 0.189 | 0.004 | 0.086 | 0.004 |
| Egger | 0.662 | 0.031 | 0.150 | 0.023 | 0.229 | 0.007 | 0.215 | 0.010 |
| **Alt: Larger sample 1, R=0.2** |  |  |  |  |  |  |  |  |
| Oracle | 1.000 | 0.000 | 0.038 | 0.012 | 0.024 | 0.001 | 0.006 | 0.000 |
| W-Egger | 0.675 | 0.030 | 0.050 | 0.014 | 0.070 | 0.003 | 0.054 | 0.003 |
| Egger | 0.354 | 0.031 | 0.071 | 0.017 | 0.083 | 0.004 | 0.190 | 0.013 |
| **Alt: Larger sample 1, R=0.5** |  |  |  |  |  |  |  |  |
| Oracle | 1.000 | 0.000 | 0.050 | 0.014 | 0.055 | 0.001 | 0.007 | 0.000 |
| W-Egger | 0.992 | 0.006 | 0.075 | 0.017 | 0.148 | 0.005 | 0.086 | 0.004 |
| Egger | 0.988 | 0.007 | 0.160 | 0.024 | 0.166 | 0.005 | 0.520 | 0.032 |
| **Alt: Larger sample 2, R=0.2** |  |  |  |  |  |  |  |  |
| Oracle | 1.000 | 0.000 | 0.058 | 0.015 | 0.065 | 0.001 | 0.015 | 0.001 |
| W-Egger | 0.500 | 0.032 | 0.079 | 0.017 | 0.077 | 0.004 | 0.052 | 0.003 |
| Egger | 0.079 | 0.017 | 0.092 | 0.019 | 0.237 | 0.017 | 0.068 | 0.004 |
| **Alt: Larger sample 2, R=0.5** |  |  |  |  |  |  |  |  |
| Oracle | 1.000 | 0.000 | 0.042 | 0.013 | 0.161 | 0.001 | 0.015 | 0.001 |
| W-Egger | 0.938 | 0.016 | 0.217 | 0.027 | 0.161 | 0.006 | 0.097 | 0.004 |
| Egger | 0.212 | 0.026 | 0.117 | 0.021 | 0.319 | 0.015 | 0.116 | 0.006 |

**Table S3:**
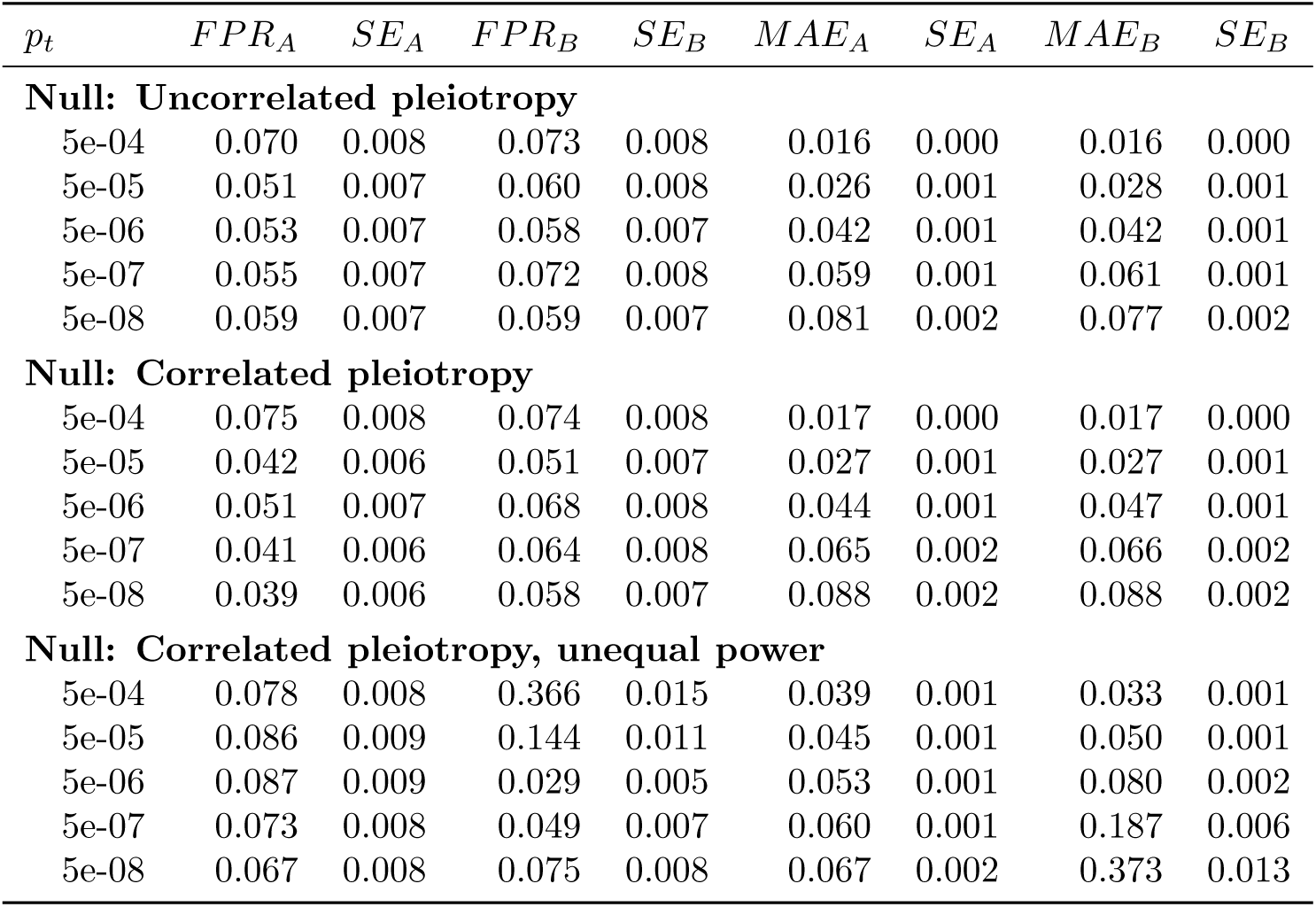
Weighted Egger regression controls the false positive rate for various *p*_*t*_ choices. We simulated GWAS summary statistics for two phenotypes (*A, B*) with *M* = 1, 000, 000 independent SNPs, 20% heritability and *N* = 100, 000 individuals in both the SNP discovery and effect estimation cohorts. In each simulation, there were 5, 000 causal SNPs per phenotype and neither phenotype had an effect on the other. When pleiotropic effects are uncorrelated, *p*_*t*_ = 5 × 10^−4^ shows a mild increase in FPR and all others control the FPR at level *α* = 0.05. The same is true when pleiotropic effects are correlated. When pleiotropic effects are correlated but there is unequal power, all cutoffs display a modest increase in the FPR for the *A* → *B* direction, and all cutoffs at or below 5 × 10^−6^ control the FPR in the *B* → *A* direction.

**Table S4:** Higher *p*_*t*_ thresholds improve power in the alt direction but increase false positives in the null direction. We simulated GWAS summary statistics for two phenotypes (*A, B*) with *M* = 1, 000, 000 independent SNPs, 20% heritability and *N* = 100, 000 individuals in both the SNP discovery and effect estimation cohorts. In each simulation, there were 5, 000 causal SNPs per phenotype and *A* has a variable effect on *B*. We find that in all settings higher *p*_*t*_ thresholds improve power in the alt direction but increase false positives in the null direction. We conclude that *p*_*t*_ = 5 × 10^−6^ greatly improves power over the standard *p*_*t*_ = 5 × 10^−8^ with only a modest increase in false positives, and an overall reduction in magnitude of the effect inferred in the reverse direction.

| $p_t$ | $PWR_A$ | $SE_A$ | $FPR_B$ | $SE_B$ | $MAE_A$ | $SE_A$ | $MAE_B$ | $SE_B$ |
| --- | --- | --- | --- | --- | --- | --- | --- | --- |
| **Alt: Equal sample sizes, R=0.2** |  |  |  |  |  |  |  |  |
| 5e-04 | 1.000 | 0.000 | 0.150 | 0.023 | 0.023 | 0.001 | 0.020 | 0.001 |
| 5e-05 | 0.988 | 0.007 | 0.079 | 0.017 | 0.048 | 0.002 | 0.031 | 0.001 |
| 5e-06 | 0.683 | 0.030 | 0.050 | 0.014 | 0.075 | 0.003 | 0.046 | 0.002 |
| 5e-07 | 0.375 | 0.031 | 0.042 | 0.013 | 0.089 | 0.004 | 0.064 | 0.003 |
| 5e-08 | 0.246 | 0.028 | 0.033 | 0.012 | 0.102 | 0.005 | 0.083 | 0.004 |
| **Alt: Equal sample sizes, R=0.5** |  |  |  |  |  |  |  |  |
| 5e-04 | 1.000 | 0.000 | 0.596 | 0.032 | 0.058 | 0.002 | 0.051 | 0.002 |
| 5e-05 | 1.000 | 0.000 | 0.288 | 0.029 | 0.108 | 0.003 | 0.065 | 0.002 |
| 5e-06 | 0.988 | 0.007 | 0.175 | 0.025 | 0.189 | 0.004 | 0.086 | 0.004 |
| 5e-07 | 0.850 | 0.023 | 0.142 | 0.023 | 0.229 | 0.006 | 0.120 | 0.006 |
| 5e-08 | 0.612 | 0.032 | 0.079 | 0.017 | 0.239 | 0.008 | 0.155 | 0.007 |
| **Alt: Larger sample 1, R=0.2** |  |  |  |  |  |  |  |  |
| 5e-04 | 0.929 | 0.017 | 0.133 | 0.022 | 0.050 | 0.002 | 0.018 | 0.001 |
| 5e-05 | 0.812 | 0.025 | 0.058 | 0.015 | 0.060 | 0.003 | 0.028 | 0.001 |
| 5e-06 | 0.675 | 0.030 | 0.050 | 0.014 | 0.070 | 0.003 | 0.054 | 0.003 |
| 5e-07 | 0.467 | 0.032 | 0.046 | 0.014 | 0.078 | 0.004 | 0.107 | 0.006 |
| 5e-08 | 0.358 | 0.031 | 0.062 | 0.016 | 0.087 | 0.004 | 0.161 | 0.010 |
| **Alt: Larger sample 1, R=0.5** |  |  |  |  |  |  |  |  |
| 5e-04 | 1.000 | 0.000 | 0.433 | 0.032 | 0.095 | 0.003 | 0.037 | 0.001 |
| 5e-05 | 1.000 | 0.000 | 0.242 | 0.028 | 0.118 | 0.004 | 0.055 | 0.002 |
| 5e-06 | 0.992 | 0.006 | 0.075 | 0.017 | 0.148 | 0.005 | 0.086 | 0.004 |
| 5e-07 | 0.958 | 0.013 | 0.085 | 0.018 | 0.157 | 0.005 | 0.185 | 0.010 |
| 5e-08 | 0.896 | 0.020 | 0.104 | 0.020 | 0.165 | 0.006 | 0.334 | 0.018 |
| **Alt: Larger sample 2, R=0.2** |  |  |  |  |  |  |  |  |
| 5e-04 | 1.000 | 0.000 | 0.092 | 0.019 | 0.039 | 0.001 | 0.041 | 0.002 |
| 5e-05 | 1.000 | 0.000 | 0.088 | 0.018 | 0.043 | 0.002 | 0.045 | 0.002 |
| 5e-06 | 0.500 | 0.032 | 0.079 | 0.017 | 0.077 | 0.004 | 0.052 | 0.003 |
| 5e-07 | 0.162 | 0.024 | 0.096 | 0.019 | 0.135 | 0.006 | 0.058 | 0.003 |
| 5e-08 | 0.080 | 0.018 | 0.088 | 0.018 | 0.216 | 0.012 | 0.066 | 0.003 |
| **Alt: Larger sample 2, R=0.5** |  |  |  |  |  |  |  |  |
| 5e-04 | 1.000 | 0.000 | 0.246 | 0.028 | 0.146 | 0.002 | 0.072 | 0.003 |
| 5e-05 | 1.000 | 0.000 | 0.212 | 0.026 | 0.098 | 0.003 | 0.078 | 0.003 |
| 5e-06 | 0.938 | 0.016 | 0.217 | 0.027 | 0.161 | 0.006 | 0.097 | 0.004 |
| 5e-07 | 0.435 | 0.032 | 0.183 | 0.025 | 0.259 | 0.011 | 0.106 | 0.005 |
| 5e-08 | 0.150 | 0.023 | 0.146 | 0.023 | 0.343 | 0.016 | 0.118 | 0.006 |

